# Changes in Protein *N*-Glycosylation Regulation Occur in the Human Parkinsonian Brain in a Region-Specific Manner

**DOI:** 10.1101/2022.05.19.492623

**Authors:** Ana Lúcia Rebelo, Richard R. Drake, Martina Marchetti-Deschmann, Radka Saldova, Abhay Pandit

**Affiliations:** CÚRAM SFI Research Centre for Medical Devices, University of Galway, Ireland; Department of Cell and Molecular Pharmacology and Experimental Therapeutics, Medical University of South Carolina, Charleston, USA; Institute of Chemical Technologies and Analytics, Vienna University of Technology, Vienna, Austria; National Institute for Bioprocessing Research and Training (NIBRT), University College Dublin, Ireland; School of Medicine, College of Health and Agricultural Science, University College Dublin, Ireland

**Keywords:** Parkinson’s Disease, human brain, N-glycosylation, protein glycosylation, glycomics

## Abstract

Parkinson’s Disease (PD) associated state of neuroinflammation due to the aggregation of aberrant proteins is widely reported. One type of post-translational modification involved in protein stability is glycosylation. Here, we aimed to characterise the human Parkinsonian nigro-striatal *N*-glycome, and related transcriptome/proteome, and its correlation with endoplasmic reticulum stress and unfolded protein response (UPR), providing a comprehensive characterisation of the PD molecular signature. Significant changes were seen upon PD: 3% increase in sialylation and 5% increase in fucosylation in both regions, and 2% increase in oligomannosylated *N*-glycans in the substantia nigra. In the latter, a decrease in the mRNA expression of sialidases and an upregulation in the UPR pathway were also seen. To show the correlation between these, we also describe an *in vitro* functional study where changes in specific glycosylation trait enzymes (inhibition of sialyltransferases) led to impairments in cell mitochondrial activity, changes in glyco-profile and upregulation in UPR pathways. This complete characterisation of the human nigro-striatal *N*-glycome provides an insight into the glycomic profile of PD through a transversal approach while combining the other PD “omics” pieces, which can potentially assist in the development of glyco-focused therapeutics.

## Introduction

Parkinson’s Disease (PD) is characterised by the death of dopaminergic neurons in the substantia nigra pars compacta due to the presence of Lewy Bodies composed of protein aggregates, leading to neurodegeneration (1). Despite the increased knowledge and pathway-oriented research regarding PD pathophysiology, the study of how glycosylation in the brain correlates with the progression of the disease remains overlooked. Glycosylation is the major highly-regulated post-translation modification of proteins, modulating the protein’s function, structure and conformational stability (2). Since PD is associated with aberrant aggregation of α-synuclein, and as glycosylation affects proteins’ structure and function, studying this feature is highly attractive.

*N*-linked glycans are the most common type of glycans in eukaryotic glycoproteins, since around 90% of these glycoproteins carry them (3). *N*-glycans can be categorized into three groups: oligomannose (only mannose residues are linked to the core structure); complex (branched structures are attached to the core); and hybrid (only mannose residues are linked to the Man-α-(1,6) arm of the core and one/two branches starting with a GlcNAc residue are on the Man-α-(1,3) arm) (4). Their ubiquity and implication in every biological process highlights their importance. In the central nervous system (CNS) they play essential roles in differentiation, synaptogenesis, neurite outgrowth and myelinogenesis during development (5)]. Congenital disorders of glycosylation (CDGs) are associated with different neuropathological symptoms such as seizures and stroke-like episodes (6), highlighting how glycan dysregulations can affect the CNS.

*N*-glycosylation takes place within the Endoplasmic Reticulum (ER) and the Golgi apparatus, involving various glycosidases and glycosyltransferases that participate in the formation and trimming of the different glycosidic chains (7,8) and chaperones to promote proper folding of the proteins. At the same time, a strict quality control system occurs by the glucosyltransferase uridine diphosphate glucose (UGGT) in the ER. This recognises any misfolded glycoproteins (which could be due to aberrant *N*-glycosylation) and either promotes their re-glycosylation until proper folding or targets them for degradation (9–11). It is also in the ER that the first step in the assembly of *N*-glycans takes place, where the newly-formed glycan chain is transferred from a lipid-linked oligosaccharide to the asparaginyl residues of the new glycoprotein’s precursors, which is catalysed by an oligosaccharyltransferase (2). The presence of metabolic deficiencies affecting this process, such as tunicamycin administration, has been described to be compensated by ER stress responses (12,13). To decrease ER stress and inhibit the accumulation of misfolded proteins, the unfolded protein response (UPR) takes place, being activated through three different signal transducers: protein kinase-like ER kinase (PERK), protein kinase inositol requiring kinase 1 (IRE1) and activation transcription factor 6 (ATF6) (14). On the other hand, (an) insufficient ER stress response has been correlated with (an) increased susceptibility of mouse cerebellar neurons to *N*-glycosylation defects (15), emphasizing the bidirectional relationship between *N*-glycosylation and ER homeostasis (16). Thus, there are multiple players whose actions must be taken into account when characterising the healthy vs diseased environment.

This study represents a comprehensive spatio-temporal analysis of the human nigro-striatal protein N-“glyco” profile to understand how it is modulated along with other molecular players in PD (Figure 1). Since glycosaminoglycans (GAGs) and O-glycome have recently been characterised in PD (17,18), we explore the missing puzzle pieces to complete the characterisation of protein glycosylation upon the onset of PD. This overview of the PD “-omics” is relevant since it allows for a more encompassing investigation of the modulation of *N*-glycosylation upon disease and how it interacts transversely with the overall molecular signature in the brain.

**Figure 1.**
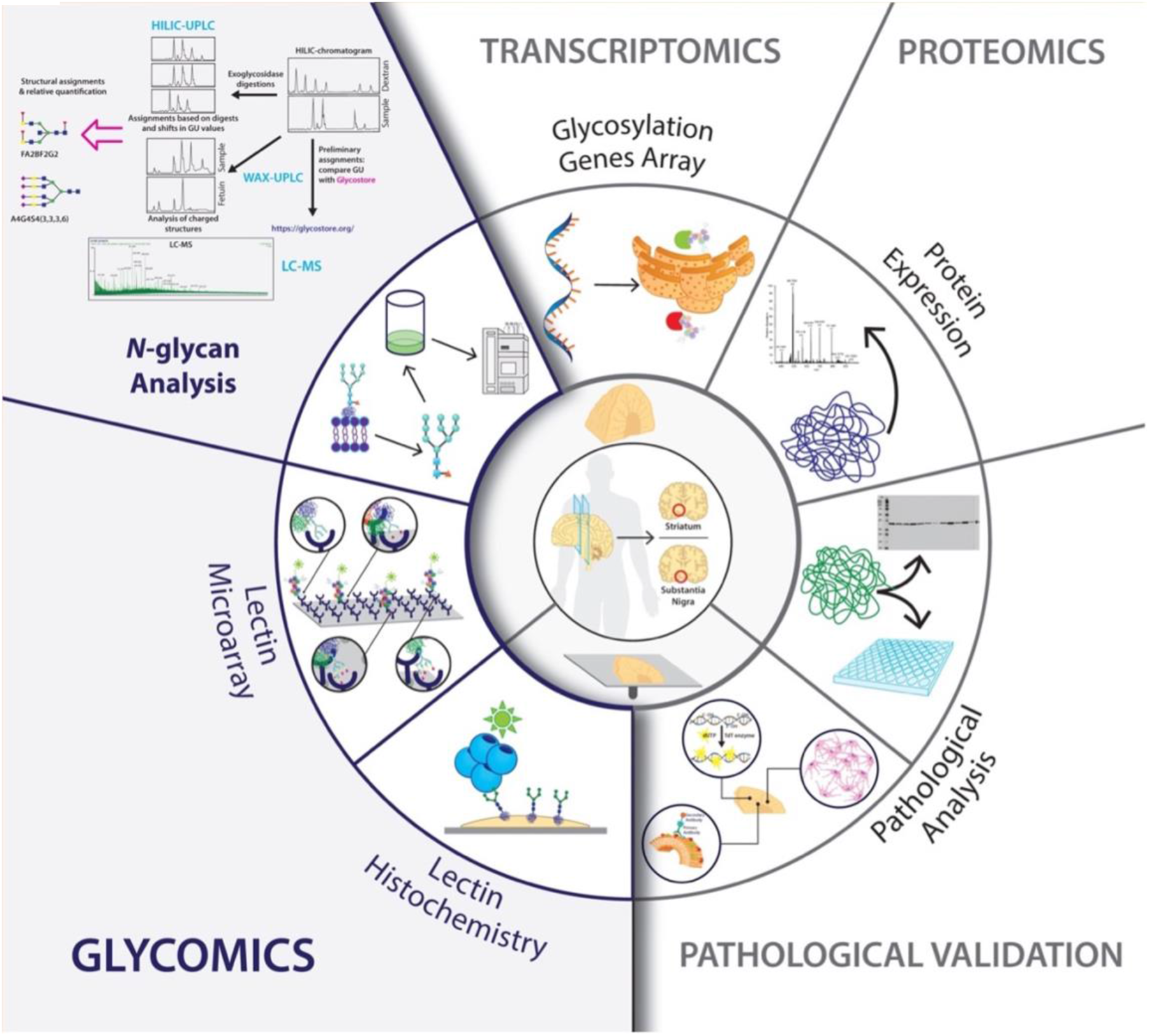
Schematic representation of the experimental design and procedures followed in this chapter. The study was designed for the region-specific and temporal characterisation of the molecular signature in the Parkinsonian brain, with a focus on *N-*glycosylation. Two regions were analysed (striatum and substantia nigra) from healthy subjects (n=18), Incidental Lewy-Body Disease (ILBD) patients (n=3) and Stage 3-4 Parkinson’s Disease (PD) patients (n=15). Brain tissue from these patients was acquired either snap-frozen or in fixed-frozen sections. For the *N*-glycome studies, a multi-faceted approach was developed using Hydrophilic Interaction Ultra Performance Liquid Chromatography (HILIC-UPLC), exoglycosidase digestions, Weak Anion Exchange Liquid Chromatography (WAX-UPLC) and Liquid Chromatography-Mass Spectrometry (LC-MS). Snap frozen tissue was used to perform glycomic, transcriptomic and proteomic analyses. Sections were used to validate the previous findings through Matrix-Assisted Laser Desorption/Ionisation Mass Spectrometry Imaging (MALDI MSI), and fluorescent and chromogenic histochemistry stainings.

## Results

### Specific glycosylation traits changes are seen in the overall glycome upon PD progression

To assess the alterations happening in the overall glycome, a high throughput screen was performed using a lectin microarray (Figure 2). A high presence of mannose residues, multi antennary *N*-glycans, GlcNAc oligomers, terminal galactose, alpha(2,6)-linked sialic acids and fucosylated glycans was detected across the groups and in both regions. Surprisingly, a low amount of Tn antigen (GalNAc--O-Ser/Thr) was also present, which is usually lacking in healthy cells but has been reported in the serum of Alzheimer’s disease (AD) patients and their age-matched controls (19). Significant differences were detected in the binding intensity of some of these lectins, which translates into a different expression of some glycans. In the striatum, a significant decrease was seen in the binding of AAA (outer arm fucose) and PWA (branching). On the other hand, in the substantia nigra there is a significant decrease in mannosylation (ASA, GNA), GlcNAc groups (DSL, LEL, PA-I), galactosylation/poly-LacNAc groups (RCA, CAL) as well as core fucosylation, which suggests that a decrease in complexity occurs. The binding of SNA and MAA did not express significant differences in disease in either of the regions.

**Figure 2.**
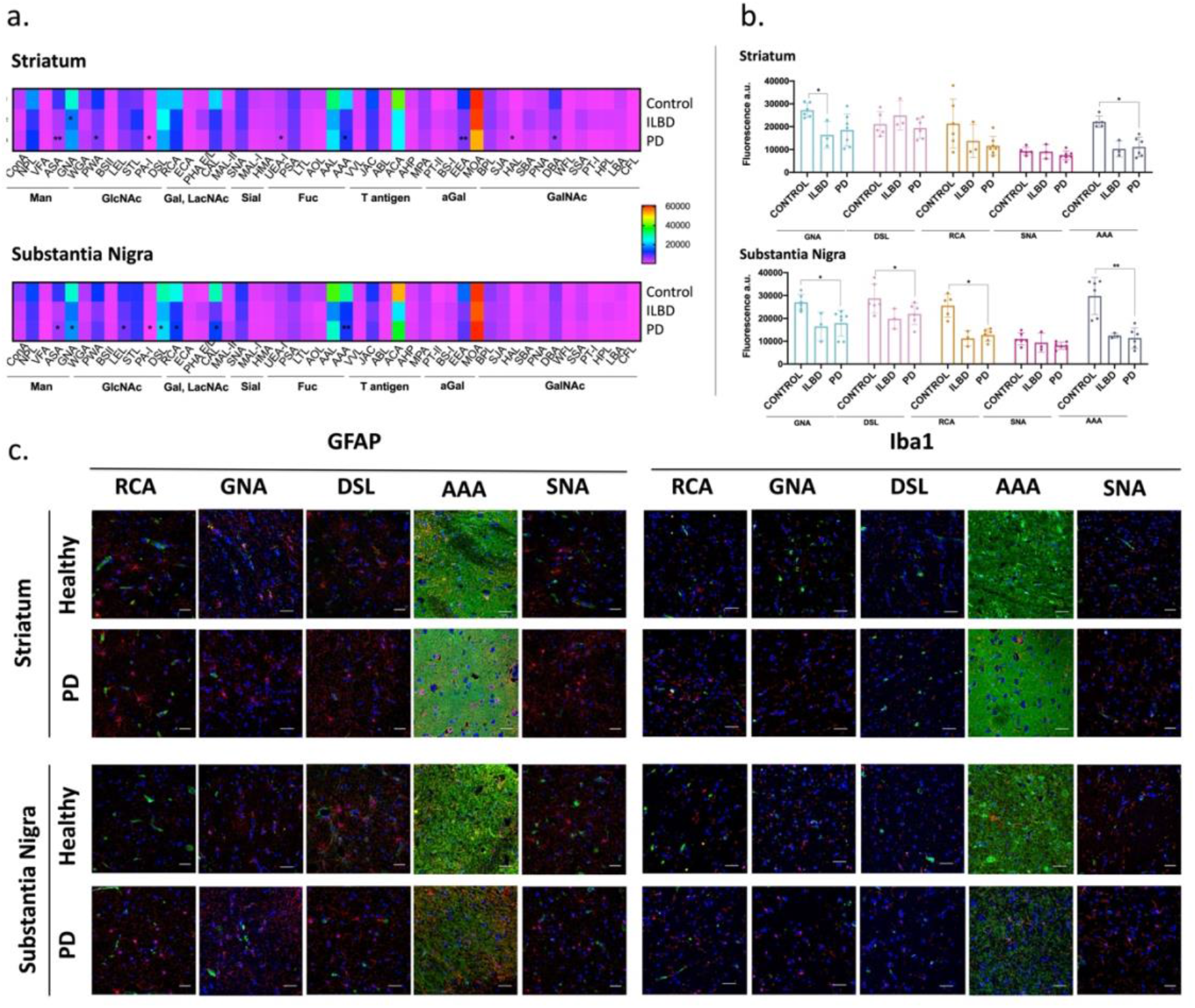
Overall changes in striatal and nigral tissue glycosylation upon Parkinson’s Disease. **a**. Lectin array assay performed on the protein lysate obtained from healthy (n=7), ILBD patients (n=3) and PD patients (n=7) to detect the expression of individual glycan structures present on both *N*- and *O*-linked glycans. Data presented as the mean ± SD. Two-way ANOVA was performed followed by Tukey’s post-hoc test and statistical significance set at *p<0.05 and **p<0.01 in relation to the healthy group. b. Relative expression of the main differently regulated glycans in the tissue (as indicated in the lectin array). c. Combined lectin and immunohistochemistry was used to study the spatial distribution of the main differently regulated glycans in the tissue (as indicated in the lectin array), and to associate their expression with specific inflammation-related cell types – astrocytes (GFAP+) and microglia (Iba1+). Scale bar = 50 μm.

Lectin histochemistry for ricinum communis agglutinin (RCA), galanthus nivalin agglutinin (GNA), datura stramonium (DSL), sambucus nigra agglutinin isolectin-I (SNA), anguilla anguilla agglutinin (AAA) combined with immunohistochemistry for the main inflammatory-related cells (astrocytes and microglia) was performed to assess the spatial distribution of the different glycans (Figure 2). Some of these correlated with astrocytes, but they were mainly binding to either extracellular matrix, or other cell types (potentially neurons).

### Parkinsonian brains display region-specific alterations in *N*-Glycosylation traits compared to the healthy ones

After looking at the overall glycosylation traits, a more in-depth study of the *N*-glycosylation patterns was performed (list of the samples used is described in Table S2; healthy controls were 1:2 male to female ratio, and the ages were 79 ± 10 years). For this, a combination of hydrophilic interaction liquid chromatography – ultra performance liquid chromatography (HILIC-UPLC) and liquid chromatography-mass spectrometry (LC-MS) was used after assessing the reproducibility of the results with control samples (Figure S1). The full *N*-glycome profiles of the healthy human nigro-striatal regions are fully described in the supplementary info (Figure S2).

The main glycosylation traits were then analysed individually to compare the *N*-glycome between healthy, ILBD and PD in both regions (according to Table S1). List of the samples used is described in Table S2: healthy controls were 1:2 male to female ratio, and the ages were 79 ± 10 years, PD patients were 1:1.2 male to female rations and ages were 80 ± 5 years). The analysis by traits rather than by individual structures is crucial since the different glyco-moieties of glycoproteins play distinct roles in facilitating the recognition, binding and processing through the different receptors on cell membranes.

In both regions the abundance of neutral glycans remained similar in healthy and incidental Lewy bodies disease (ILBD), decreasing significantly in the later stages of the disease (Figure 3). Polysialic acids were significantly increased by 1.6-fold in the striatum upon disease; however, the opposite trend was seen in the substantia nigra (11% reduction).

**Figure 3.**
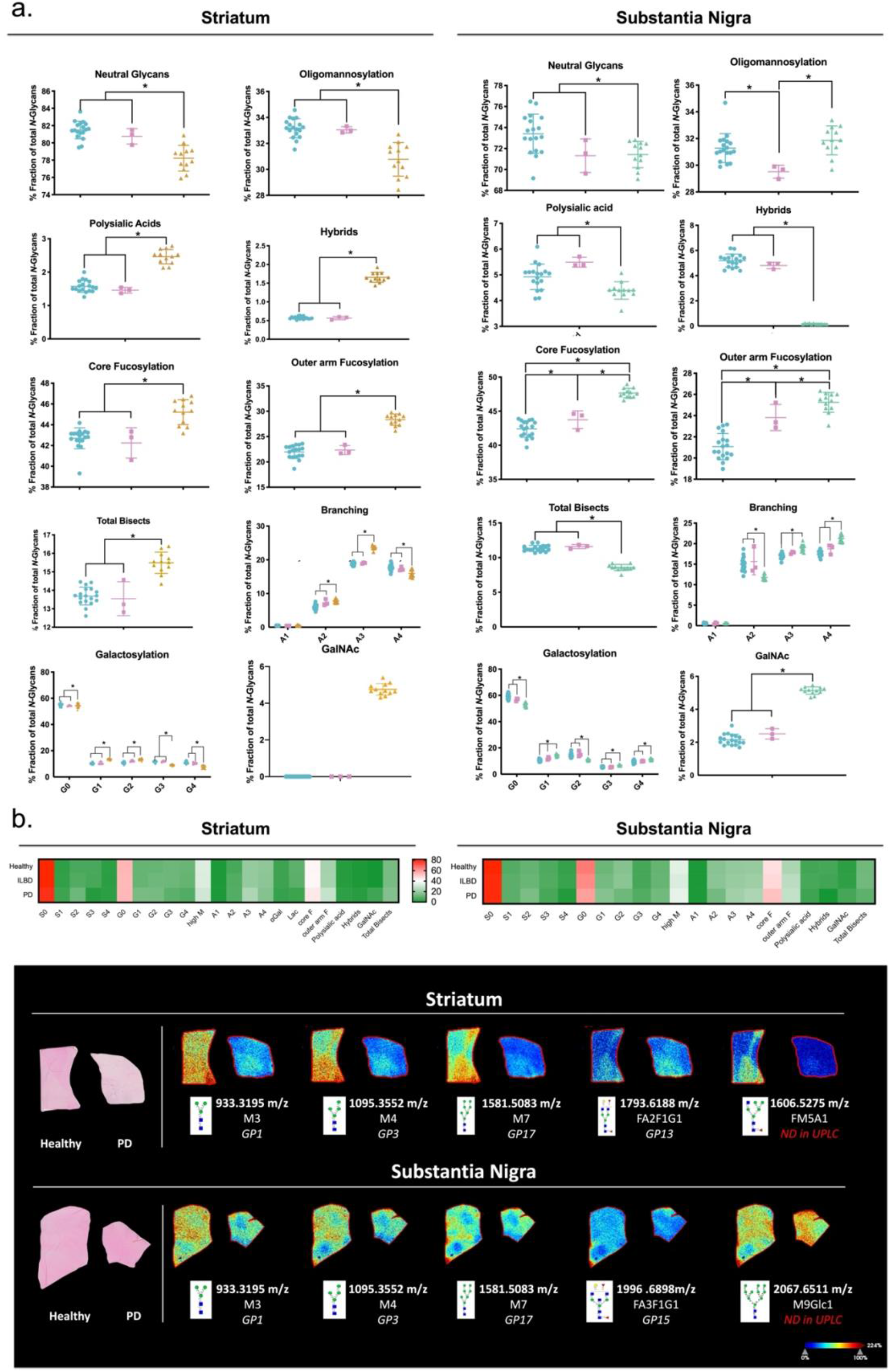
Changes in the *N*-glycosylation traits in the different regions, in healthy and upon disease. a. The common glycosylation features amongst the main structures in each glycan peak were grouped in the main glycosylation traits (according to Table 2.5). The abundance of these was log transformed. Data is presented as mean ± SD. One-way ANOVA was performed since the data was shown to be normally distributed. This was followed by Tukey’s post-hoc test and statistical significance set at *p<0.05. Blue: healthy; Pink: ILBD; Yellow/Green: PD. b. Differences seen in terms of abundance amongst the main glycosylation traits. c. *N*-glycan imaging of the main structures differently regulated in the striatum and substantia nigra of PD and age-matched control human brains. Each image is accompanied by the putative structures determined by combinations of accurate m/z, CID fragmentation patterns and glycan database structure. ND – non-detected

Concerning oligomannosylation, in the two regions analysed the trends seen were contrasting: in the striatum the abundance of oligomannose structures seems constant between healthy and the ILBD group, decreasing significantly at the later stages of the disease. However, in the substantia nigra there is a decrease in the presence of these glycans upon ILBD, and an increase at the later stages of the disease (Figure 3).

As for fucosylation, in both regions a significant increase in both core and outer arm fucosylation between healthy controls and later stages of PD was seen (in the striatum, 1.09-fold increase in core fucose and 1.28-fold increase in outer arm fucose, whereas in the substantia nigra 1.12-fold increase in core fucose and 1.19-fold increase in outer arm fucose were seen). In the substantia nigra there is a significant increase in both types of fucosylation in the ILBD group, which suggests that fucose expression occurs in parallel with the progression of the disease. On the other hand, the expression of galactose residues in the *N*-glycan structures present both in the striatum and substantia nigra increases significantly upon PD (1.01-fold and 1.11-fold, respectively). Interestingly, there is a significant increase in mono-galactosylated structures in both regions (1.30-fold and 1.38-fold respectively), while the tri- and tetra-galactosylated ones decrease in the striatum (21% and 32% reduction, respectively) and increase in the substantia nigra (1.28-fold and 1.20-fold respectively).

Regarding the branched structures, *N*-glycans with two branches are significantly increased in the striatum (1.31-fold increase), whereas they decrease in the substantia nigra (22% reduction). However, glycans with three branches are upregulated in both regions. Structures with four antennae are sharply decreased in the striatum and increased in the substantia nigra. Also, there are no significant changes between the healthy and ILBD groups in either the bisected or the branched glycans.

Individual *N-*glycans’ identification, validation, and spatial distribution were assessed through MALDI mass spectrometry imaging (MALDI-MSI). This revealed a decreased expression of specific structures in PD brains compared to healthy ones, corroborating the significant differences already determined by HILIC-UPLC. This was the case of M3, M4, M5 and FA2F1G1 (which correspond to GP1, GP3, GP17 and GP13, respectively, on the HILIC-UPLC data) in the striatum (Figure 3c). Another structure (FM5A1) was shown to be downregulated in PD brains through MALDI-MSI; however, this was not detected through HILIC-UPLC. This contributes to the decrease in oligomannosylation seen upon disease in the striatum. Similarly, in the case of the substantia nigra, downregulation in the expression of some oligomannose structures was also seen, mainly of M3, M4 and M7, corresponding to GP1, GP3 and GP17, respectively (Figure 3c). A decrease in FA3F1G1 (GP15) is also seen through MALDI-MSI, and the presence of another structure (M9Glc1 – increased upon PD), which was not previously detected through HILIC-UPLC. This indicates that even though there is an overall increase in oligomannosylation in the substantia nigra upon disease, the abundance of some specific structures from this glycosylation trait is decreased. Other structures were detected; however, significant changes were not seen using this technique.

The results seen highlight the possibility of using this technique not only to validate the previous data, but also to assess its spatial distribution within the different regions in the striatum and substantia nigra. Furthermore, this also allows the previous data to be complemented through the detection of structures that were not deciphered previously by HILIC-UPLC and LC-MS.

### Regulation of glyco-enzymes transcripts and expression is region- and disease-dependent

As previously mentioned, glycosylation of proteins is a non-template-driven mechanism that depends upon the action of multiple glyco-enzymes (2). Therefore, an assessment of the changes in their transcription and expression is of significance.

In the substantia nigra of PD patients, significant downregulation was mainly seen in N-acetylgalactosaminyltransferases (GANTL-5, -9, -11, -14, which are predominantly involved in O-glycosylation, were decreased by -14.65, -3.64, -3.91 and -8.59-fold, respectively), in N-acetylglucosaminyltransferases (MGAT-4B, -4C, -5B, which play a role in the branching of *N*-glycans), in sialidases (NEU-1, -2) and sialyltransferases (ST8SIA-3, -4, -6, which are mainly regulating the formation of poly-sialic acids). There was also a -3.75-fold significant decrease in GLB1, a galactosidase involved in the different types of protein glycosylation and in the processing of gangliosides, more specifically GM1, which is reported to be reduced in PD brains (20)(Figure 4).

**Figure 4.**
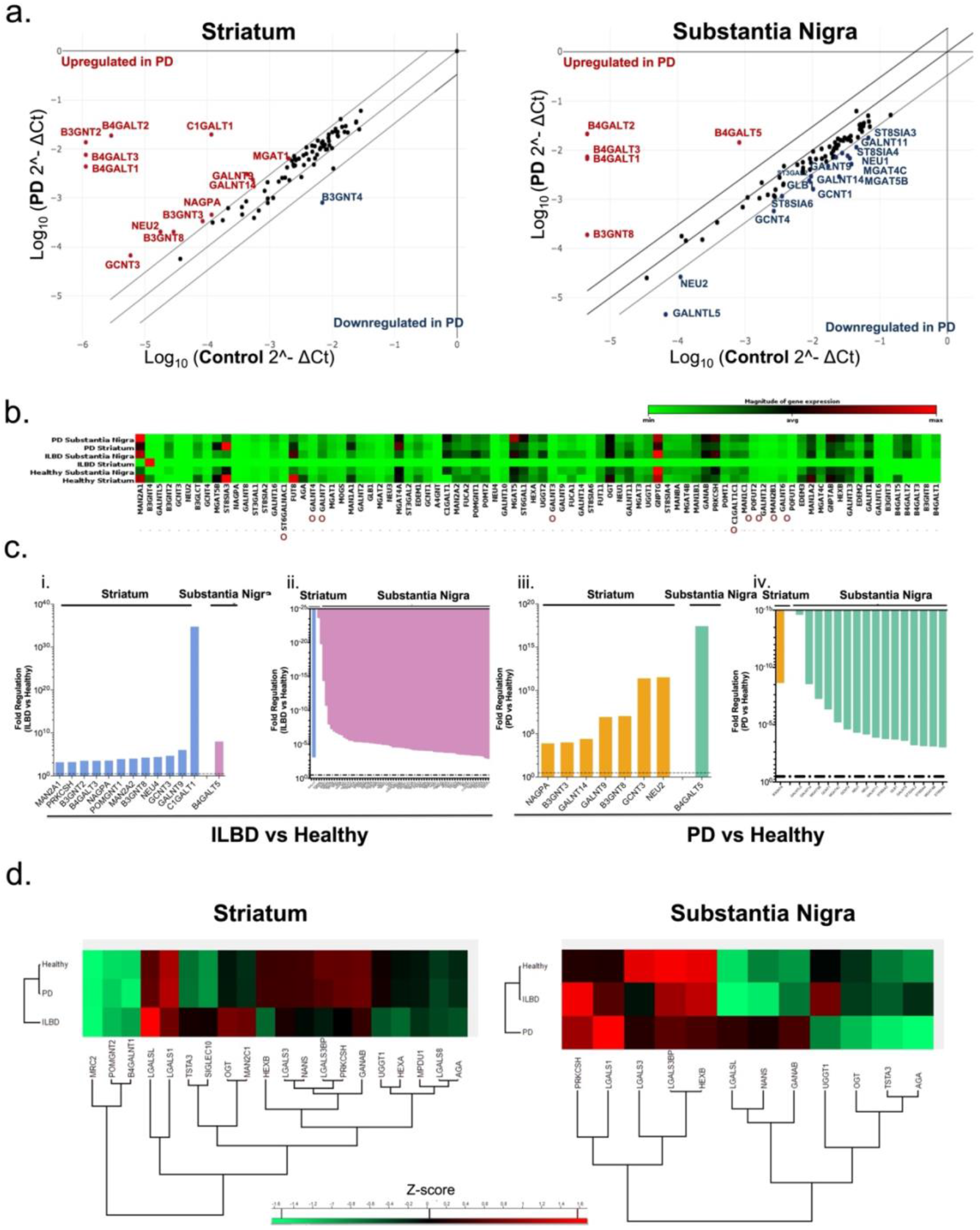
Expression of the human nigro-striatal glycoenzymes upon Incidental Lewy Bodies Disease and Parkinson’s Disease. **a**. Expression of 84 transcripts of glycosylation enzymes was performed using RT^2^ Profiler™ PCR Array - Human Glycosylation (Qiagen®) and analysed through the GeneGlobe Data Analysis Center®. Significant upregulation (indicated in red) or downregulation (indicated in blue) of the different genes was considered when fold change ≥ 3 and p ≤ 0.05. **b**. Relative expression of each transcript in healthy, ILBD and PD samples. Red indicates high relative expression and green low relative expression. Expression levels were normalised on the house keeping gene GAPDH. **O** indicated enzymes involved only in *O*-glycosylation. All the others are involved only in *N*-glycosylation or in both types of glycosylation. **c**. Genes significantly upregulated (i., iii.) or downregulated (ii., iv.) in ILBD (i., ii.) and in PD (iii., iv.) conditions, in both the striatum and substantia nigra (considered when fold change ≥ 3 . Data represents the pool of mRNA from six biological replicates for the healthy group, three for ILBD group and six for PD group. **d**. Relative expression of the glycoenzymes detected by proteomics (through nanoLC-MS) in health, ILBD and PD samples according to their z-score (red: upregulated proteins, and green: downregulated proteins).

In the striatum, only one gene in PD was significantly downregulated - B3GNT4 (-8.69-fold reduction), an N-acetylglucosaminyltransferase responsible for the biosynthesis of poly-N-acetyllactosamine sequences. In contrast, there were multiple genes whose transcription was upregulated in this region: two N-acetylglucosaminyltransferases (B3GNT-3, -8, related to the elongation of branched *N*-glycans), two N-acetylgalactosyltransferases (GALNT-9, -14, involved in the formation of O-glycan core structures) and GCNT3, also involved in O-glycosylation (Figure 4). The expression of a sialidase (NEU2) is also significantly increased (by 11.51-fold).

In the case of the striatum, in the ILBD group there is a significant upregulation in mannosidases expression (3.14-fold and 4.00-fold increase in MAN2A-1, -2, respectively, which are the final players in the *N*-glycans maturation pathway, controlling the conversion of the oligomannose structures to complex ones), and enzymes involved in O-glycosylation (C1GALT1, GALNT9, GCNT3, POMGNT1) (Figure 4). It is also important to mention the significant upregulation of NAGPA (3.52-fold) and B3GNT-2, -8, seen both in the ILBD and PD striatum, which might indicate that the changes in these pathways start from the early stages of the disease. The expression of protein kinase C substrate 80K (PRKCSH) transcript is significantly increased as well in ILBD. This is a crucial enzyme for the formation of *N*-glycans since it cleaves glucose residues from the LLO formed in the *N*-glycans synthesis. Interestingly, multiple enzymes responsible for regulating the initial stages of the *N*-glycans formation are upregulated in the ILBD group. This might suggest that an adaptive response from the cellular machinery is taking place to compensate for the initial toxicity seen with the formation of the Lewy bodies. However, it is worth keeping in mind that the expression of the genes does not entirely reflect the expression of the final form of the enzymes. Also, their activity might be affected due to the pathophysiological conditions of the cellular milieu.

To assess the expression of the enzymes whose transcripts were evaluated previously, the proteomic profile was assessed by nanoLC-MS. Unfortunately, the expression of the glyco-enzymes of interest is low, so only very few of them were detected (Figure 4).

### Increase in specific ER stress markers is seen in the Substantia Nigra of PD brains

Since the ER plays a pivotal role in regulating protein homeostasis and is closely related to *N*-glycosylation, UPR was also assessed through the expression of different markers using western blot. There were no significant changes in the markers assessed in the striatum, apart from the expression of PDI, which was significantly downregulated upon PD (31.3% reduction) (Figure 5). In the case of the substantia nigra, there was a significant increase in the expression of partial ATF6 (2.38-fold increase) and PDI (1.51-fold increase)-classic markers of UPR activation (Figure 5).The expression of other chaperones (GRP78, GRP94) was not significantly altered upon disease in either of the regions studied using western blot. However, when the proteomic analysis was performed using nanoLC-MS, an upregulation was found in these chaperones and PDI in the substantia nigra, indicating an overall upregulation in this canonical pathway (Figure 5). The undetected changes seen through western blot can be due to the defective binding between the antibody and the antigen present in each of these proteins.

**Figure 5.**
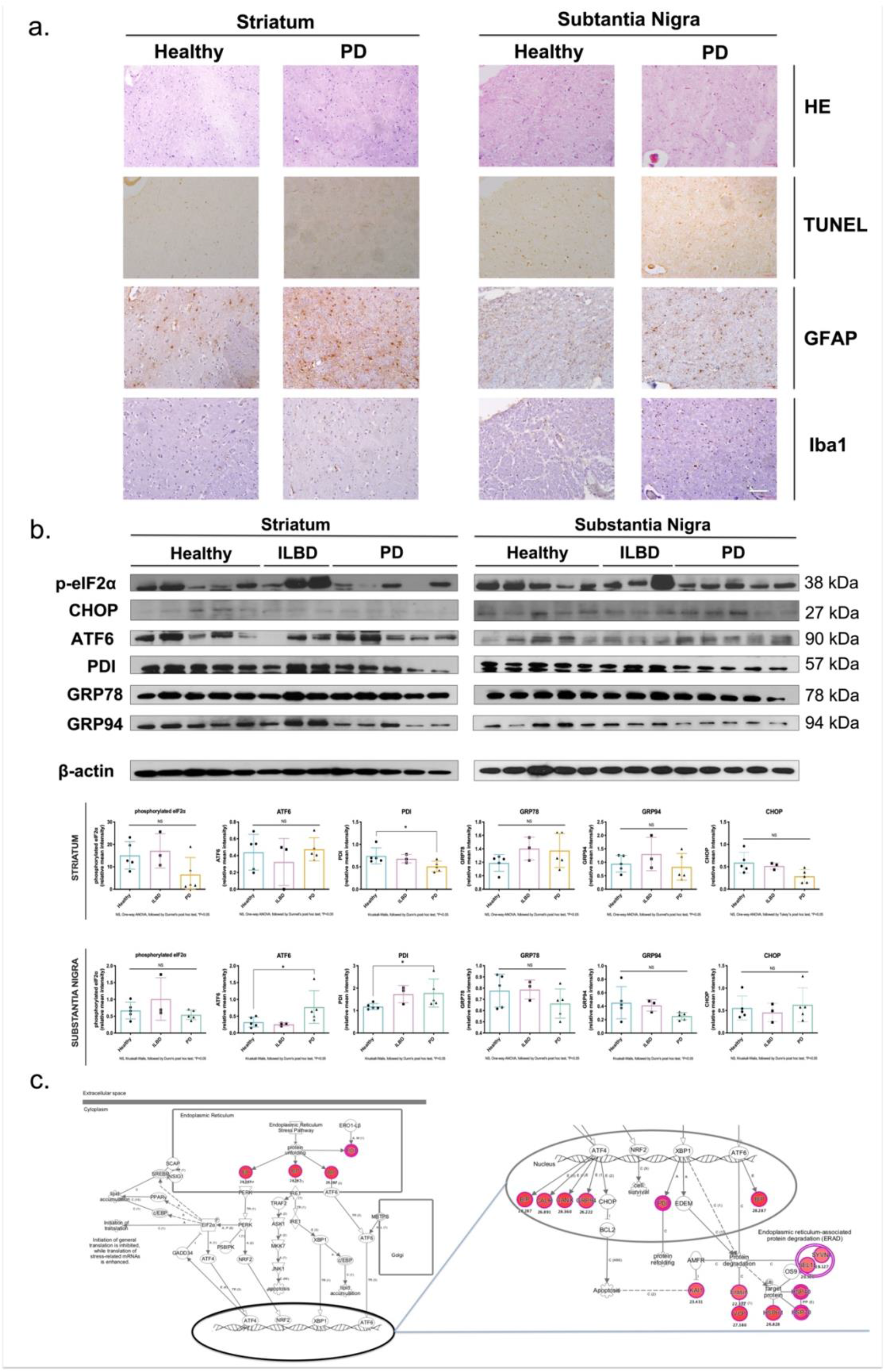
Pathological markers regulation upon PD in the striatum and substantia nigra with a focus on the endoplasmic reticulum stress (ER stress) and unfolded protein response (UPR). a. Morphological characterisation and spatial analysis of the distribution of the main neuroinflammation-related cells in both the striatum and substantia nigra upon PD. (i) Heamatoxylin/Eosin (H&E) staining, (ii) terminal deoxynucleotidyltransferase dUTP nick end labelling (TUNEL) staining, (iii) anti-GFAP (astrocytes marker) immunohistochemistry, (iv) anti-Iba1 (microglia marker) immunohistochemistry. Scale bar = 100 μm. b. Expression of ER stress/UPR related markers (GRP78, GRP94, PDI, ATF6, phosphorylated eIF2a, CHOP) was assessed and quantified through western blot analysis. If the data was normally distributed, one-way ANOVA was performed and followed by Dunnett post-hoc test, and statistical significance set at *p<0.05. If the data was not normally distributed, Kruskall-Wallis test followed by Dunn’s post hoc test were carried out, and statistical significance was set at *p<0.05. c. Ingenuity Pathway Analysis (IPA) of canonical ‘unfolded protein response’ in Parkinsonian substantia nigra, based on the proteomic analysis performed after running the proteins in nanoLC-MS. Red symbols indicate identified and upregulated activation of proteins in the signalling pathway from a diseased brain associated with ER stress.

### Downregulation in the expression of sialyltransferases leads to impairments in cell metabolism, changes in glyco-profile and increase in ER stress (Proof-of-concept *in vitro* correlation study)

The characterization of the nigro-striatal *N*-glycome, the regulation of the glyco-enzymes involved in such phenomenon and the changes in the ER homeostasis provided in the previous paragraphs describe important information regarding the glyco-profiling of PD human brains. However, it is difficult to understand whether glycosylation is acting as a cause or an effect in the onset of neurodegenerative cascades. In fact, it is highly likely that it acts as both, promoting a cyclic chain of events. It was already reported that neuroinflammation leads to changes in glycosylation (27, 52). Nonetheless, the opposite is yet to be demonstrated. Deciphering this can be a dauting task, so we carried out a small *in vitro* functional study as a “proof of concept” highlighting how a change in glycosylation can lead to other molecular alterations in the PD context. Moreover, we emphasize how all of these molecular events are connected and could potentially be targeted in future therapies.

As glycosylation is a very complex and sophisticated phenomenon, for this study, the only glycosylation trait considered was the decrease in poly-sialylation in the substantia nigra, as well as the decrease in sialyltransferases (ST8SIA-3, -4, -6 – mainly involving the formation of poly-sialic acids) seen upon disease in this brain region. To assess whether it is the downregulation of these enzymes that could be leading to the changes in the glycome, cell homeostasis, and ER stress, a mixed primary cell culture model using rat embryonic cells from the ventral mesencephalon (origin of dopaminergic neurons) was used, and a glycosyltransferase inhibitor (more specifically a sialyltransferase inhibitor) was employed at different concentrations (50 μM, 250 μM and 500 μM). After 48h of incubation, it was seen that the inhibition of these enzymes (sialyltransferases) led to changes in the expression of different glycans (decrease in sialylation and increase in mannosylation), in the mitochondrial activity of the cells (showing that neuronal degeneration is occurring) (in a dose-response fashion) and also in the dysregulation in the expression of UPR genes (increase in the mRNA expression of XBP1 spliced, ATF6 and PDI) at the highest concentration (Figure 6). An increase in inflammation was also seen upon the decrease in sialylation, as is shown by the increase in GFAP expression (reactive astrocytes).This emphasizes the impact that glycosylation can have in diseased states and how it can lead to a cascade of molecular events.

**Figure 6.**
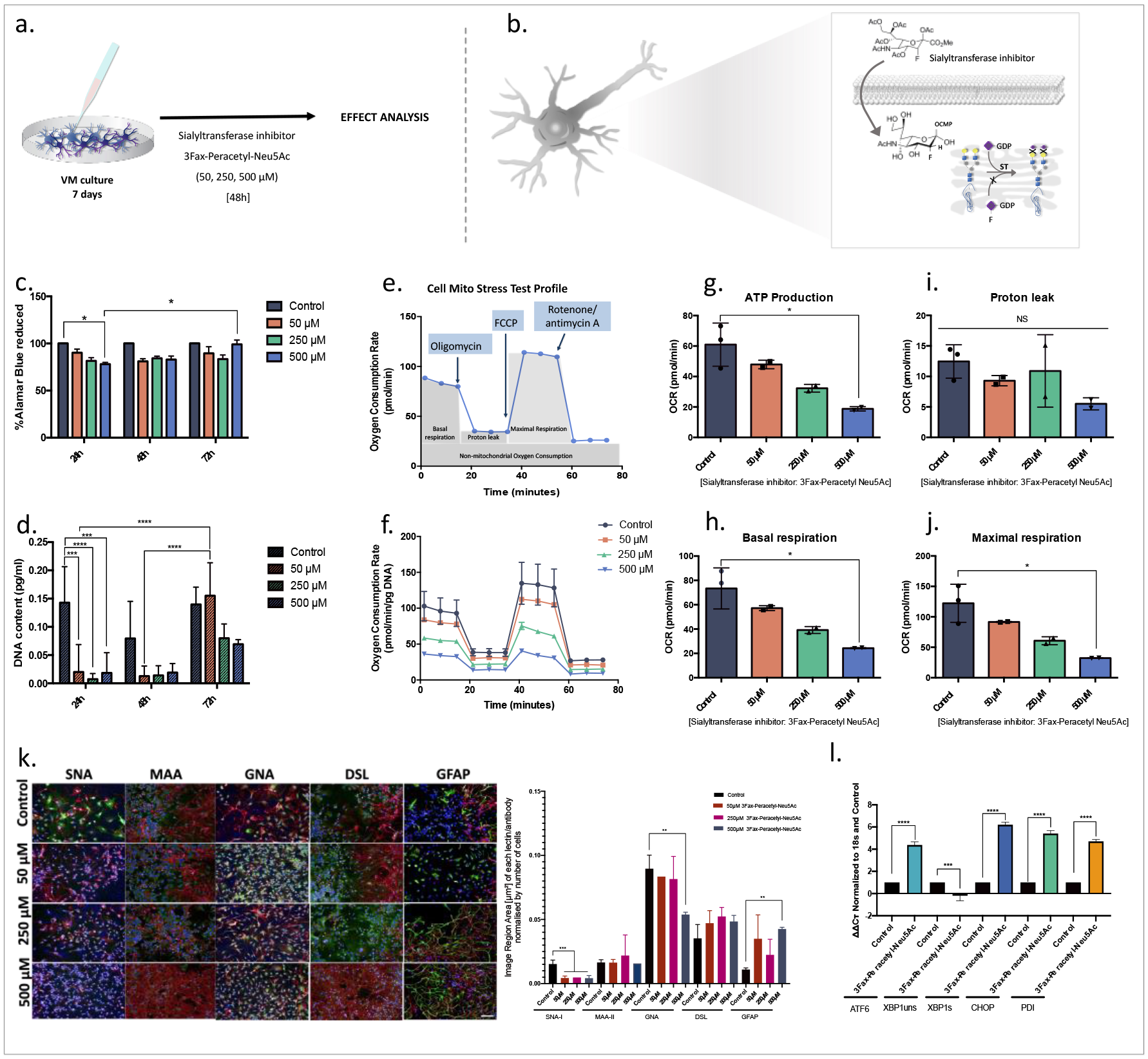
*In vitro* correlation study: Influence of the downregulation of sialyltransferases in the cell metabolism, glyco-profile and ER stress. a. Schematic outline of the *in vitro* model, time points and concentrations of 3Fax-Peracetyl Neu5Ac used; b. Mechanism of action of 3Fax-Peracetyl Neu5Ac – it crosses the cell membrane and inhibits the attachment of sialic acid residues to glycan structures in the Golgi apparatus; c. Metabolic activity of VM cells when in contact with 3Fax-Peracetyl Neu5Ac assessed through alamarBlue assay (n = 3, data presented as the mean ± SD). Two-way ANOVA was performed, followed by Tukey post-hoc test, and the statistical significance was set at *p<0.05, **p<0.01, ***p<0.001; d. Cell proliferation of VM cells when in contact with 3Fax-Peracetyl Neu5Ac studied through PicoGreen assay (n = 3, data presented as the mean±SD). Two-way ANOVA was performed, followed by Tukey post-hoc test, and the statistical significance was set at *p<0.05, **p<0.01, ***p<0.001; e.-j. Mitochondrial activity of VM cells when in contact with 3Fax-Peracetyl Neu5Ac assessed through SeaHorse® Cell Mito Stress Test ® (Agilent, USA). In e. the standard profile in a SeaHorse® test is described, and in f. the oxygen consumption rate (OCR) profile after each compound is added to the media is described (function of each compound is described in the materials and methods). In g. – j. the ATP production, proton leak, basal respiration and maximal respiration are described for the VM cells when in contact with different concentrations of 3Fax-Peracetyl Neu5Ac (n = 3, data presented as the mean ± SD). One way ANOVA, followed by Tukey’s post hoc and statistical significance was set at *p<0.05. It shows that only the highest concentration of 3Fax-Peracetyl Neu5Ac decreases significantly the cells ATP production, basal respiration and maximal respiration; k. Combined lectin (in green) and immunohistochemistry (BIII-Tubulin antibody in red) was used to study the expression of the main types of glycans in the VM cells upon contact with different concentrations of 3Fax-Peracetyl Neu5Ac at 48h, and to associate their expression with specific neurons (BIII-Tub+). Scale bar: 100 μm, One-way ANOVA was performed followed by Holm-Sidak’s post-hoc test and the statistical significance was set at **p<0.01, ***p<0.001 – A strong decrease in SNA-I binding is seen (sialylation) as well as in GNA (mannosylation), there is also an increase in GFAP (astrocytes expression), showing an upregulation in inflammation; l. mRNA expression of different UPR players is described in controls (VM cells alone with vehicle) and cells incubated with the highest concentration of 3Fax-Peracetyl Neu5Ac (500 μM). ATF6, XBP1s, CHOP and PDI show significant increase upon decrease in sialylation, One way ANOVA, followed by Tukey’s post hoc and statistical significance was set at *p<0.05, **p<0.01, ***p<0.001.

## Discussion

The focus of this study was to characterise the protein glycosylation profile upon disease by studying the changes in the different *N*-glycosylation traits, which complements the “glyco-” studies already published on the dysregulation of GAGs (17) and the O-glycome (18) in human brain PD samples. *N*-glycosylation traits were analysed in parallel and correlated with the alterations seen in the transcriptomics and proteomic expression of glyco-enzymes and the UPR in the two regions studied and in the two diseased conditions (ILBD and PD Braak stages 3/4) (overall results seen are summarised in Figure 7).

**Figure 7.**
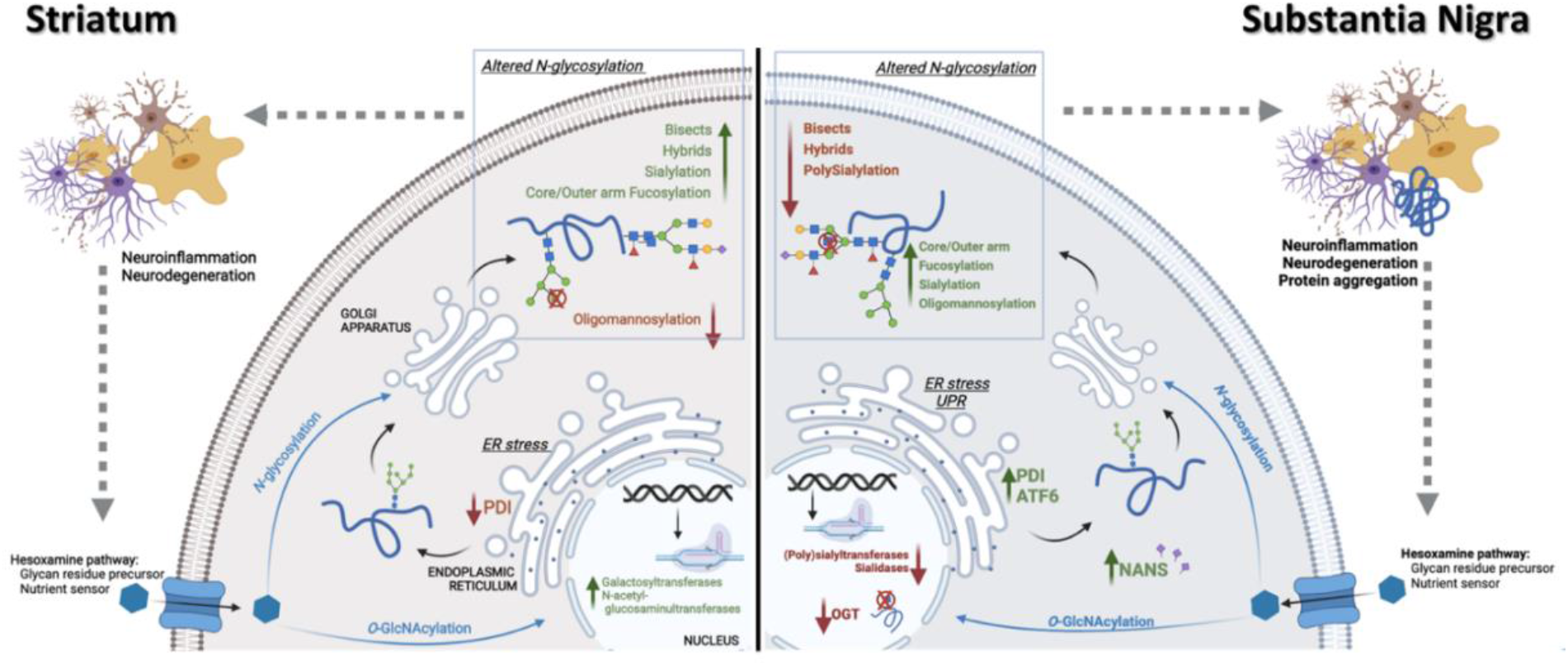
Summary and main conclusions seen regarding the molecular signature in each brain region upon PD. Differential regulation of glycosylation traits, glycosylation enzymes and UPR-related proteins is seen in the striatum vs substantia nigra. Created with BioRender®.

The overall high abundance of fucosylated and oligomannosylated structures seen through the lectin array corroborates what was reported previously (21,22). This array also showed the presence of GalNAc-containing O-glycans, being in accordance with the study exploring the O-glycome of PD brains (18). Some significant changes were seen in the substantia nigra, where a decrease in the expression of lectins binding to mannose, galactose and GlcNAc was detected upon PD. This is a methodology which allows for high-throughput analysis, being of use as a preliminaty assessment of glycomic alterations. However, it is worth noting that lectins have a broad range of ligands, so they can be used to detect carbohydrates in both glycoproteins and glycolipids, failing to show any distinction in the nature of the glycosylated molecule (23). There are also issues related with the accuracy of quantification and the possibility of unspecific binding leading to noise signal. Because the focus of this study was to elucidate specifically the *N*-glycome of the human brain, further analyses were performed using an optimised multi-faceted approach combining liquid chromatography, exoglycosidase digestions and mass spectrometry, which allowed elucidation of the in-depth composition of the nigrostriatal *N*-glycoprofile, and then investigation of the alterations in each glycosylation trait upon disease.

An overall low degree of sialylation seen (around 20%) accords with previous studies reporting the *N*-glycome from other human brain regions (21) and a study on the *N*-glyco profile of the nigrostriatal pathway in the rat (24). The increase in sialic acid seen in the *N-*glycans from both regions upon PD occurs in parallel with the increase in sialylation in the O-glycans from the same regions (18). This is in contrast to studies on the *N*-glycome profiling of IgG from PD patients where sialylation is reduced (25). Increases in sialylation have been associated with brain tumours (26) to escape from immune checkpoints, which suggests that there are proteins constituents of Lewy Bodies that are oversialylated, preventing their elimination. However, such hypothesis will need to be further studied. A decrease in sialylation has been associated with neuroinflammatory models, mainly through the upregulation of neuraminidase 1 and 4 (NEU1 and NEU4), which constitute an essential factor for pathophysiological consequences upon inflammation (4,27). Similar changes in the expression of neuraminidases are described in this study as an increase in the expression of NEU2 and NEU4 transcripts in the striatum upon disease is seen. However, the overall sialylation is increased, raising the question as to whether this is due to the action of sialyltransferases or to impairments in the lysosomal degradation of such proteins (28). On the other hand, in the substantia nigra the expression of NANS was significantly upregulated, indicating a higher abundance in sialic acid substrates upon PD. Additionally, the downregulation of NEU1 and NEU2 transcripts upon PD is involved with the overall increase of sialylation. However, such conclusions will require further analysis of these enzymes’ expression since they were not detected through proteomics.

Polysialic acids (PSA) are known to be involved in synaptic plasticity in brain development and were reported to be upregulated in the hippocampal regions of AD patients with neuronal loss and aggregation of amyloid plaques (29). The decrease seen in the PSA expression in the substantia nigra is parallel to decreased expression of the sialyltransferases ST8SIA3, ST8SIA4 and ST8SIA6 transcripts, suggesting the role of these sialyltransferases on the assembly of PSA.

Oligomannose structures are usually transient biosynthetic structures of N-linked glycans that in the case of the CNS are transported to the cell membrane to integrate the cellular glycocalyx as part of recognition molecules like NCAM or an adhesion molecule on glia (AMOG) (5). In the adult brain, these are mainly present at the synapses, being the extent and quality of mannosylation regulated with synaptic maturation. Additionally, expression of mannose-binding lectins (MBL) has been reported in the brain, predominantly in astrocytes and microglia which are chief participants in controlling immune responses (30) through the lectin pathway of complement activation. An increase in mannose structures abundance suggests an increased binding to these receptors, which leads to immune cell activation and inflammatory response. PD is associated with a chronic state of inflammation and reports have shown that the expression of cytokines varies between striatum and substantia nigra, amongst healthy and ILBD and PD patients (30), enhancing the inflammatory pathology. This is further confirmed by the high expression of Iba1+ and changes in microglial morphology in our study. Additionally, since these structures are abundantly present at the synapsis, and synaptic disruption was recently reported in the PD substantia nigra through PET imaging of synaptic vessel glycoprotein (31), one can hypothesise that these oligomannose residues-carrying proteins are incorrectly released to the extracellular space, that are recognised as foreign and binding to the MBL, which activates the complement system.

In the case of the striatum, it seems that the decrease in mannose expression accompanies the decrease in dopamine. It has been shown that dopaminergic agonists that act on adenylate cyclase-linked dopamine receptor sites induce a dose-dependent enhancement in the incorporation of mannose into glycoproteins in ex vivo models of rodent striatal cultures (32), so it can be hypothesised that a reversed effect is seen due to the depletion of dopamine in this brain region. Moreover, there is a significant increase in the expression of some mannosidases, which can trim the oligomannose structures and reduce their abundance in the diseased group.

Concerning fucosylation, its importance in *N*-glycans in the CNS requires further study. Fucose residues can be added to the glycan structures by the activity of one or more of the thirteen fucosyltransferases reported in the human genome; however, only nine are involved in the formation of *N*-glycan structures (33). None of these enzymes was detected through proteomic analysis, nor were their transcripts differently regulated, so it was impossible to conclude if their activity was involved with the detected changes in fucosylation. Models that fail to express core fucosylation (FUT8-knock out) show an impairment in hippocampal long-term potentiation, presenting a schizophrenia-like behaviour and phenotype (34). This is accompanied by a significant increase in the abundance of microglia and astrocytes and their susceptibility to pro-inflammatory compounds, reaffirming that these phenotypes are due to defective neuronal physiology and dysfunctional glia populations (35). Considering this, a decrease in core fucosylation in the PD brain would be expected. But conversely in this study, an upregulation was seen. This is likely a contradiction in the understanding of the neurodegeneration pathway of PD that mediates pathfinding for neurons or direct neurite migration (33). However, this hypothesis will have to be further assessed.

As for galactose residues, these are particularly important in the healthy brain since sialylated *N*-glycans with multiple β1,3-linked galactoses and repeating units of PolyLac and a terminal galactose are abundant (36). Also, galactose residues that belong to N-acetyllactosamine moieties are recognised by galectins (e.g. galectin-1, -3, -9), which have reportedly been expressed by activated microglia and astrocytes, and are involved in neuromodulation in CNS pathophysiology, mainly by controlling inflammatory processes (37). Therefore, the increase in galactosylation seen was expected. Proteomic analysis showed a slight upregulation in galectin-1 expression in the substantia nigra but no changes were seen in the striatum.

The abundance of branched *N-*glycans is low; however, they are an essential posttranslational modification for neuronal survival since the knock-down of N-acetylglucosaminyltransferase 1 (MGAT1; enzyme responsible for initiating the formation of hybrid glycans) leads to apoptosis and severe behaviour impairments, tremors and premature death (38). The abundance of bisecting and branched *N*-glycans is intimately connected since both involve adding GlcNAc residues by N-acetylglucosaminyltransferases (MGAT) to a mannose donor, and the existence of bisecting GlcNAc can limit the synthesis of branched structures. The presence of the bisecting GlcNAc (mediated by MGAT3) inhibits the action of other glycosyltransferases (mainly MGAT4, MGAT5) (39). The expression of MGAT3 was described to increase in the AD human brain, reflecting an adaptive response to protect the brain as this increase occurs post-accumulation of β–amyloid and reduces Aβ production to protect against neurological degeneration (39). Even though the expression of MGAT3 transcript was not altered in PD brains, it is possible that the expression and/or activity of the enzyme is altered, so the increase in bisected glycans seen in the striatum can be perceived as being adaptive and protective.

The dysregulation of certain *N*-glycan structures was also reported through MALDI-MSI in AD brains (40). A region-specific modulation of *N*-glycans upon disease was also seen in this case. For example, FA2G1 (1647 m/z), FA2G2 (1809 m/z) and FA3G1 (1850 m/z) were all shown to be decreased upon AD in the hippocampus, but increased in the cortex from the diseased brains (40). Although these facts were noteworthy, the protective or adaptative role upon disease of these structures remains to be elucidated. Interestingly, FMA5A1 (1606 m/z, decreased in the PD striatum) was also decreased in the hippocampus of the AD brains. On the other hand, FA2F1G1 and FA3F1G1 (1793 m/z and 1996 m/z, respectively) were both increased in the cortex of AD brains, whereas they were downregulated in the striatum and substantia nigra (respectively) of PD brains. This indicates that these might be specific to each condition and potentially play different roles on the onset of distinct neurodegenerative diseases, also depending on the brain region where they are expressed.

The current study did not allow the tracking of the proteins in which the glycans are being modified nor the corresponding *N*-glycosylation sites where such alterations occur. This would require a glycoproteomic approach, which is only recently starting to be explored. This can provide important insights in the future, once such technologies are further developed (reviewed by (54, 55)). Nonetheless, the type of analysis carried out in the current study is pioneer in proving the full N-glycome profile in the human striatum and substantia nigra, which can already guide and help to funnel the potential future studies to be performed using glycoproteomics strategies.

It is also worth observing that single-cell glycomic analysis would be of utmost interest since it would allow for a specific investigation into in which type of cells these changes are occurring and would help to further elucidate the mechanisms occurring. However, this is still highly complex since the technologies available do not reach such resolution and sensitivity. For instance, the limited spatial resolution of current MALDI-MSI systems translates into the acquisition of spectra that encompasses molecular information from multiple adjacent cells, preventing cellular resolution. Nonethelss, the recent development of new MALDI systems with an oversampling approach is coupled with laser post-ionisation (MALDI-2) is being optimized, and was tested preliminarily in brain tissue, which might be the ideal strategy for follow up studies (56).

Concerning the interplay between ER stress and changes in *N-*glycosylation, it has been reported that the increase in spliced X-box binding protein 1 (XBP1) due to the activation of the IRE-1 branch of the UPR leads to a decrease in bisecting glycans and sialylation and increases oligomannose structures (41), which corresponds to the nigral PD *N*-glycomic signature. Here we saw changes in the ATF6 branch of the UPR and in other markers in the substantia nigra, which indicate that the UPR plays a role in the alterations seen in the *N*-glycomic profile since polysialylation and bisects decreased significantly and mannosylation increased in this region.

It was shown that ER stress is linked with dopaminergic neuronal death as different markers were shown to be significantly upregulated post-stimuli with toxins such as 6-OHDA or MPP+, both *in vitro* (42) and *in vivo* (43). Also, UPR activation was correlated with the aggregation of α-synuclein in human samples (44). Nonetheless, most of these studies cover only a few of the players involved in UPR. Therefore, it is essential to look at the overall picture and assess the different cascades of the UPR response, mainly the ATF6 pathway, which hitherto has been overlooked. Only some of the markers were differently regulated through western blot (PDI and ATF6) in this study. However, this analysis was further complemented by proteomic analysis, where an upregulation of chaperones was seen (BiP and GRP94). The main function of PDI is to catalyse the formation and rearrangement of disulphide bonds in molecules (essential for proper protein folding). It has been reported that PDI has anti-inflammatory properties, inhibiting the LPS produced by macrophages (45). Therefore, the increase in the expression of PDI seen in the substantia nigra is in line with the inflammatory profile seen and confirms an increase in UPR. Since there is also the accumulation of aggregated α-syn in these samples (demographic data supplied by Parkinson’s UK Brain Bank), it confirms the link between UPR and protein aggregation (44).

The upregulation in the expression of the cleaved ATF6 in the substantia nigra also indicates the activation of another branch of the UPR. Cleavage of ATF6 was not detected through proteomics, so western blot was the best approach to assess this. This cleavage also leads to the upregulation of PDI, which explains the increase in both of these markers seen in the substantia nigra of PD patients (46). In a rodent model of PD (MPP+) it was shown that ATF6 was significantly increased post injury, and that animals deficient in ATF6 displayed increased loss of dopaminergic neurons (47). It is, possible, therefore, that the changes seen in the different situations relate to the dynamic and flexible nature of the UPR and/or to the techniques used to quantify these changes. In either case, it is essential to explore the different cascades and players involved in the UPR.

The physiological glycome is extremely complex and sophisticated, making it hard to fully comprehend its function and, more specifically, the role of each glycosylation trait. Thus, most studies on the functional role of glycosylation in the brain were done using gene inactivation models, where specific glycosylation enzymes (commonly glycosyltransferases) were knocked-out in *in vivo* models, to evaluate the influence that the lack of a particular glycan residue would have on neuronal physiology (34). However, in the past decades the use of inhibitors for these enzymes (instead inhibiting them genetically) has emerged as a potential pharmacological tool to not only explore the biological functions of these glycosylation traits, but also to be used as possible therapeutics targeting the glycome. All the above-mentioned data characterizes changes seen in the different molecular pieces of the PD puzzle; nonetheless, it is still hard to properly correlated them. As an attempt to understand further, and since the nigro-striatal glycome is also very complex and a plethora of players were seen to be dysregulated, we opted to choose one specific glycosylation trait and test *in vitro* how an alteration in this trait this would correlate with the other molecular mechanisms that we have reported above. By using a sialyltransferase inhibitor (3Fax-Peracetyl Neu5Ac) and a culture of embryonic cells which are the cells that lead to the maturation of dopaminergic neurons (main cell type affected in PD substantia nigra), it was possible to clearly highlight how a decrease in the action of sialyltransferases (and subsequent sialylation), also affected mannosylation (increasing it), promoted inflammation (increase in GFAP+ expression, which is a marker for astrocytic reactivity), affected the mitochondrial activity of these cells (promoting neurodegeneration) and also led to an increase in UPR markers expression. All of these individual alterations seen, were also the ones we described previously to be seen in the human substantia nigra upon PD. This small “proof of concept” study emphasizes that the changes in the expression glycosylation enzymes seen could be the ones promoting the other molecular alterations. Nonetheless, it is note-worthy that this is just a small example taken from one glycosylation trait. Understandably, there are many other traits involved and a clear understanding of which are the upstream vs downstream players in the overall signature of PD is still far from being clarified.

To sum up, this study presents for the first time a comprehensive overview of the different “omic” branches of PD that relate to *N*-glycosylation in a region-dependent manner, providing information on how they interplay with disease progression, exploring pathways that were previously overlooked (Figure 7). While it is naive to assume that these dysregulations are directly correlated, it is still a valid starting point that was not considered to date. Having this overview of how the *N*-glycome is altered in parallel with the other pieces of the PD molecular signature such as the expression of glyco-enzymes, the UPR and the proteomic profile can help to preliminarily establish a pattern in a non-template-driven process. Further studies on how ER stress in the dopaminergic circuitry might be leading to *N*-glycosylation dysregulation and the investigation of single-cell glycoproteomics will be of utmost interest.

## Supporting information

Suppp file

## Acknowledgments

The authors would like to acknowledge the Parkinson’s UK Brain Bank, funded by Parkinson’s UK, a charity registered in England and Wales (258197) and in Scotland (SC037554) for supplying the tissue samples and associated clinical and neuropathological data. The authors are grateful to Mr Anthony Sloan and Dr Raghvendra Bohara for proof-reading this manuscript and Mr Maciej Doczyk for designing Figure 1. The authors are thankful to Ms Paula Kenny (National Institute for Bioprocessing Research and Training, Dublin, Ireland) for her help in performing the DMB analysis. The authors thank Mr Stefan Kirnbauer (Technical University of Vienna, Austria) for the help while establishing the method to acquire *N*-glycan detection through MALDI-MSI. The authors also acknowledge the facilities and scientific and technical assistance of the Centre for Microscopy & Imaging at the National University of Ireland Galway, facilities that are funded by NUI Galway, and the Irish Government’s Programme for Research in Third-Level Institutions, Cycles 4 and 5, National Development Plan 2007– 2013 (www.imaging.nuigalway.ie). Finally, the authors are thankful to DC Proteomics (UK) for performing the proteomic analytical service and to Asparia Glycomics (San Sebastian, Spain) for providing the lectin microarray analytical service.

This publication has emanated from research supported by a research grant from Science Foundation Ireland (SFI), co-funded under the European Regional Development Fund through Grant number 13/RC/2073 and 13/RC/2073_2, and by the BrainMatTrain project, which is funded by the European Union Horizon 2020 Programme (H2020-MSCA-ITN-2015) under the Marie Skłodowska-Curie Initial Training Network and Grant Agreement No. 676408.

## Author Contributions

A.L.R. performed all the experiments except the lectin array. R.R.D. developed the method for MALDI imaging of N-glycans. M.M.D. supervised the N-glycan characterisation and analysis through MALDI imaging. R.S. managed and supervised the N-glycan characterisation and analysis through liquid chromatography and mass spectrometry. A.P. supervised, managed the overall study and secured funding. A.L.R. wrote the manuscript, which was edited and approved by all co-authors.

## Materials and Methods

The study was designed for the region-specific and temporal characterisation of the molecular signature in the Parkinsonian brain, with a focus on *N-*glycosylation. Two regions were analysed (striatum and substantia nigra) from healthy subjects (n=18), Incidental Lewy-Body Disease (ILBD) patients (n=3) and Stage 3-4 Parkinson’s Disease (PD) patients (n=15). Brain tissue from these patients was acquired either snap-frozen or in fixed-frozen sections (Figure 1).

### Human brain tissue

Frozen autopsied striatum and substantia nigra from patients with Parkinson’s Disease (n=12 (snap frozen tissue), n=6 (fixed-frozen, 10μm thick sections)) or Incidental Lewy Bodies Disease (n=3 (snap frozen tissue), n=1 (fixed-frozen, 10μm thick sections)), healthy matched controls (n=15 (snap frozen tissue), n=6 (fixed-frozen, 10μm thick sections)), and associated clinical and neuropathological data were supplied by the Parkinson’s UK Brain Bank, funded by Parkinson’s UK, a charity registered in England and Wales (258197) and in Scotland (SC037554). The gender, age, post-mortem delay, duration of the disease and Braak stages are described in Table S2. All experiments were done in accordance with the NUI Galway Research Ethic Committee guidelines, under the reference of 18-Mar-20.

### Tissue homogenisation

The snap frozen brain tissue was homogenised in different ways depending on the assay performed. For the glycomic analysis, the snap frozen brain tissue was homogenised in RIPA® buffer (R0278, Sigma, Ireland) and cOmpleteTM Protease Inhibitor Cocktail (5056489001, Roche, Ireland, 1:25) through mechanical disruption using Qiagen TissueLyser LT (Qiagen, UK), at 4oC (40Hz, 8 minutes). The homogenates were centrifuged at 16.000 g for 20 minutes at 4oC and the supernatants were collected for further analysis. The protein concentration of the supernatants was calculated using the PierceTM BCA Protein assay kit (23225, Thermo-Fisher, UK). For the proteomic analysis, tissue was lysed in 100mM Tris (pH 8.5) with 1% sodium deoxycholate, 10 mM TCEP, 40 mM chloroacetamide and cOmpleteTM Protease Inhibitor Cocktail. The samples were vortexed, boiled for 5 minutes and sonicated on ice. The homogenates were centrifuged at 20.000 g for ten minutes and the supernatants were collected for further analysis.

### *N*-glycan analysis

### Isolation and fluorescence labelling of *N*-glycans

The isolated glycoproteins were dried in a vacuum centrifuge overnight (Savant™ SPD131DDA SpeedVac™ Concentrator, ThermoFisher). These were immobilized in an acrylamide gel and reduced and alkylated. *N*-glycans were released using *N*-glycanase PNGase F (1239U/ml, New England BioLabs, Inc. cat no. P0709L) and were fluorescently labelled with 2-aminobenzamide (2-AB) by reductive amination (24,49,50), and the excess of 2-AB was removed on WhatmanTM 3MM paper (Clifton, NJ) in acetonitrile washes (51).

Hydrophilic interaction liquid chromatography – ultra performance liquid chromatography (HILIC-UPLC) UPLC was performed using a UPLC Glycan BEH Amide Column, 130A, 1.7 μm particles, 2.1 x 150mm (Waters, Milford, MA) on an H Class AcquityTM UPLC (Waters, Milford, MA) equipped with a Waters temperature control module and a Waters AcquityTM fluorescence detector. Solvent A was 50 mM ammonium formate, pH4.4 and Solvent B was acetonitrile. The column temperature was set to 40oC and the sample temperature to 5oC. The method had a duration of 30 minutes and it was composed of a linear gradient of 30% to 47% of buffer A for 24 minutes at 0.561mL/min flow rate, increasing to 70% at minute 25 and returning to 30% at minute 27 until the end of the run. Samples were injected in 70% acetonitrile. Samples were excited at 330 nm and fluorescence was measured at 420. nm. For each sample set, the system was calibrated using a dextran ladder of 2AB-labelled glucose oligomers (Waters, Milford, MA), as described elsewhere (50).

Weak anion exchange – ultra performance liquid chromatography (WAX-UPLC) WAX-UPLC was carried out using a DEAE anion exchange column, 10 μm particle size, 75 x 7.5mm on a Water AcquityTM UPLC separations module complete with a Waters AcquityTM UPLC fluorescence detector. Solvent A consisted of 20% v/v acetonitrile in water and solvent B was 0.1M ammonium acetate buffer, pH7.0, in 20% v/v acetonitrile. This method had a duration of 30 minutes and the gradient conditions were a linear gradient of solvent A from minute 5 to minute 20 from 100% to 0%. At minute 23 solvent A returns to 100%, remaining constant until the end of the run, all at a 0.750 mL/min flow rate. Samples were injected into water. Samples were excited at 330 nm and fluorescence was measured at 420 nm. For each sample set, the system was calibrated using *N-*glycan extracted from fetuin (50).

### Matrix-assisted laser desorption/ionisation mass spectrometry imaging (MALDI-MSI) of *N-*glycans

For spatial *N*-glycome analysis through MALDI mass spectrometry imaging, samples were prepared as previously described (52). Further info on sample preparation can be found on the Supplementary methods. Released *N*-glycan ions were detected using a MALDI FT-ICR scimaXTM (Bruker Daltonics, Germany) operating in positive mode with a Smart Beam II laser operating at 1000 Hz and a laser spot size of 20 μm. Signal was collected at a raster width of 150 μm between spots. A total of 300 laser shots was collected to form each pixel. Following the acquisition, data was processed and images of expressed glycans were generated using FlexImagingTM 5.0 and SCiLSTM Lab 2021 software (Bruker Daltonics, Germany), where ions in the range of 900-3200 m/z were analysed. Observed mass/charge ratios (*N*-glycans) were searched against the glycans assigned and detected through LC-MS previously, glycan databases using GlycoWorkbench® and the database provided by the Consortium for Functional Glycomics (www.functionalglycomics.org). Represented glycan structures were also generated in GlycoWorkbench®, as they were determined by a combination of their measured accurate m/z, CID fragmentation patterns and previous structural characterisation carried out by HILIC-UPLC.

### Gene array

Brain tissue from the striatum and substantia nigra was homogenized and lysed in TRI Reagent® (Sigma, Ireland). The RNA was extracted from this, and its concentration, purity and integrity were checked using the NanoDropTM 2000c Spectrophotometer (Thermo Scientific, UK) (reading at 260/230 nm and 260/280 nm of wave length), and the BioAnalyzerTM 2100 (Agilent, USA). cDNA synthesis was performed using the RT2 first strand kit (Qiagen, UK) and following the manufacturer’s instructions.

cDNA was labelled with SyBR Green using an RT2 SyBR Green Mastermix provided by the manufacturer, which contained a HotStartDNA® Taq Polymerase (Qiagen, UK). This mixture (containing the labelled cDNA) was dispensed into the 384-wells of each plate (10 uL per well) containing 86 primers for the genes encoding for glycosylation genes such as glycosidases and glycosyltransferases (Table S5). The plate was inserted in the Roche LightCycler® 480 (Roche Life Science, Switzerland) and the cycles were defined as follows: 1 cycle for ten minutes at 95oC, and 45 cycles of 15 seconds at 95oC and one minute at 60oC. The data acquired was analysed using the GeneGlobe Data Analysis Centre® (Qiagen, UK). Significant upregulation or downregulation of the different genes was considered when fold change ≥ 3 and p ≤ 0.05. The collected data represents the pool of mRNA from six biological replicates for the healthy group, three for the ILBD group and six for the PD group.

### Proteomic analysis

Proteins (200 μg) were digested with trypsin at an enzyme to substrate ratio of 1:50 (w/w) for 18 hours at 37 oC, and then stopped by acidification with formic acid (FA). to a final concentration of 2% (v:v). Peptides were desalted using C18 Sep-Pak cartridges following the manufacturer’s instructions, dried, and the peptide concentration was determined using the Thermo Scientific™ Pierce™ Quantitative Fluorimetric Peptide Assay. Peptides were reconstituted in 100 mM Triethylamonium bicarbonate (TEAB), and tandem mass tag (TMT) labelling was carried out on 30 μg of peptides from each individual sample. Samples were distributed in the 11-plex label set. TMT labelling reagents were each dissolved in anhydrous acetonitrile. Each sample containing 30 μg of peptide in TEAB buffer was combined with its respective 11-plex TMT reagent and incubated for one hour at room temperature. Then, 5% hydroxylamine was added, and the combined sample incubated for a further 15 minutes. This mixture was then dried for further analysis.

#### Nano-LC mass spectrometry

The dried sample was resuspended in 5% formic acid and desalted using a SepPak® cartridge according to the manufacturer’s instructions (Waters, UK). Eluate from the SepPak® cartridge was again dried and resuspended in buffer A (20 mM ammonium hydroxide, pH 10) prior to fractionation by high pH reversed-phase chromatography using an UltiMate® 3000 liquid chromatography system (Thermo Scientific). For reversed-phase chromatography, an XBridge BEHTM C18 Column (130 Å, 3.5 μm, 2.1 mm X 150 mm, Waters, UK) was used, and peptides were eluted with an increasing gradient of buffer B (20 mM Ammonium Hydroxide in acetonitrile, pH 10) from 0% to 95% over 60 minutes. The resulting fractions were dried and resuspended in 1% formic acid prior to analysis by nano-LC MSMS using an Orbitrap Fusion Tribrid mass spectrometer (Thermo Scientific). Here, peptides in 1% (vol/vol) formic acid were injected onto an Acclaim PepMap® C18 nano-trap column (Thermo Scientific). After washing with 0.5% (vol/vol) acetonitrile 0.1% (vol/vol) formic acid, peptides were separated on a 250 mm x 75 μm Acclaim PepMap® C18 reverse phase analytical column (Thermo Scientific) over a 150 minutes organic gradient. Solvent A was 0.1% formic acid and Solvent B was aqueous 80% acetonitrile in 0.1% formic acid. Organic gradient included 7 gradient segments (1% to 6% of solvent B over one minute, 6% to 15% of solvent B over 58 minutes, 15% to 32% of solvent B over 58 minutes, 32% to 40% of solvent B over five minutes, 40% to 90% of solvent B over one minute, and then kept at 90% of solvent B for six minutes and then reduced to 1% of solvent B over one minute) with a flow rate of 300 nL/min. Peptides were ionized by nano-electrospray ionization at 2.0kV using a stainless steel emitter with an internal diameter of 30 μm (Thermo Scientific) and a capillary temperature of 275 oC. All spectra were acquired using an Orbitrap FusionTM TribridTM mass spectrometer controlled by Xcalibur® 2.0 software (Thermo Scientific) and operated in data-dependent acquisition mode using an SPS-MS3 workflow. The raw files were analysed with MaxQuant® v. 1.6.17 and searched against the latest human fasta database (version 05.2020, 20368 sequences) downloaded from UniProt. Identified peptides and proteins were filtered to 1% FDR. TMT reagent impurity correction was performed.

#### Data analysis: Perseus workflow and Ingenuity pathway analysis (IPA)

MaxQuant® data was exported to Perseus® software for complete proteomics analysis, where the results were log2 transformed as well as filtered to exclude any contaminants and non-significant peptides. The final data set (including the fold ratio change of each group compared to healthy and p-values) was further analysed in the Ingenuity Pathway Analysis software (IPA®, Qiagen, UK), where canonical pathways and upstream regulator data were compared by their activation z-score and associated p-value. The main proteins and pathways considered related to unfolded protein response, glyco-related proteins/enzymes and neuroinflammatory signaling.

### *In vitro* cell culture model

#### Embryonic ventral mesencephalon rat cells extraction

Ventral mesencephalic (VM) cells were isolated from day 14 embryonic (E14) Sprague-Dawley rats. Summarily, pregnant Sprague-Dawley rats were euthanised by decapitation under isofluorane, the yolk sac was removed by laparotomy, the embryos excised from the sac and the brains separated from their heads on ice-cold Hank’s Balanced Salt Solution (HBSS) (Sigma, Ireland). The mesencephalon was removed from each brain, and the VM tissue was cut from this. The tissue was trypsinized, manually dissociated and suspended in media, and the isolated cells were incubated in a humidified atmosphere of 5% CO2 at 37 oC and cultured in media (Dulbecco’s modified Eagle’s medium/F12, 1% penicillin-streptomycin (PS), 1% fetal bovine serum (FBS), 33 x 10-3 M D-glucose, 1%L-glutamine, supplemented with 2% B27). All the reagents mentioned are from Sigma, Ireland, except B27, which is from ThermoFisher, Ireland. Each experiment was done in triplicate.

#### Sialylation-deficient model

To assess the effect of sialylation on neuronal/glial physiology, a fluorinated sialyltransferase inhibitor was put in contact with VM cell cultures. This inhibitor competes actively with sialic acids (Neu5Ac/Neu5Gc), preventing the action of sialyltransferases in including sialic acids in glycan structures during their processing in the Golgi. After seven days in culture, 3Fax-Peracetyl-Neu5Ac (566224, Merck, Ireland) was incubated with the cells at 50, 250 or 500 μM for 48 hours in a humidified atmosphere of 5% CO_2_ at 37 oC. Subsequently, the cells were collected and the effects of the inhibitor on cell physiology were analysed. Further methods are described in the supplementary materials and methods.

### Biological assays *in vitro*

To evaluate the effects of the sialyltransferase inhibitor on the physiology of the cellular culture, multiple assays were carried out, as described below.

#### Cell metabolic activity: AlamarBlue®

To investigate the cells metabolic activity at different concentrations of sialyltransferase inhibitor, alamarBlue® assay (Invitrogen, Ireland) was used. Briefly, at the pre-defined time-points, the cells were incubated with a 10% solution of alamarBlue in HBSS for three hours. Subsequently, the absorbance from the alamarBlue solution was read at 450 nm and 550 nm using a Varioskan Flash plate reader (ThermoFisher, UK). HBSS and alamarBlue (100%) alone were used as negative and positive controls, respectively, for the succeeding analysis.

#### DNA quantification: Quant-iT PicoGreenTM dsDNA assay

Quant-iTTM PicoGreenTM dsDNA assay kit (Invitrogen, Ireland) was used to assess the DNA content (direct measure of cell proliferation). To collect the samples, the media was removed from the cells, these were washed with HBSS, and then DNase free water was added to each well. These underwent three cycles of freezing-thawing (alternating between at least 15 minutes at -80

°C and at least 1 hour at room temperature). Afterwards, samples were incubated with TE buffer (at 1X) and PicoGreen reagent (at 1X) for ten minutes in a 96-well plate (protected from light) and the fluorescence was read at 520 nm using a Varioskan Flash plate reader (ThermoFisher, UK).

#### Mitochondrial activity: MitoStress test (Seahorse extracellular flux analysis system®)

To study the cultures’ mitochondrial activity in real time, more specifically the oxygen consumption rate (OCR) and associated cellular basal and maximal respiration, ATP production and proton (H+) leak, the Seahorse extracellular flux analysis system and corresponding MitoStress test (Agilent Technologies, USA) were used as per manufacturer’s instructions (and following the method already published (57)). Briefly, cells were seeded at 2.5 x 10^4^ cells/well into 8-well Seahorse XFp cell culture miniplates (Agilent Technologies, USA) and treated with the sialyltransferase inhibitor for 48 hours. Then, the culture medium was replaced with Seahorse assay medium (Seahorse Base XF medium, with 10 mM D-glucose, 2 mM L-glutamine, 1mM Na+ pyruvate, at pH 7.4) and the cells were left for one hour prior to the assay in an incubator without CO_2_ at 37 ºC. Mitochondrial function was investigated through bioenergetics profiles that were generated by sequential injection of multiple mitochondrial toxins resuspended in Seahorse assay medium: 1) oligomycin A (an ATP synthesase inhibitor that blocks its proton channel/complex V), 2) carbonyl cyanide-4- (trifluoromethoxy)phenylhydrazone (FCCP; an uncoupling agent that disrupts the mitochondrial membrane potential by transporting hydrogen protons through the membrane and leading the respiration rate to its maximum capacity), and 3) rotenone/antimycin A (complex I and III inhibitors, respectively, which shut down mitochondrial respiration). These were loaded in the cartridge in a way to reach the final optimised concentrations of 1.5 μM, 3 μM and 0.5 μM, respectively, in the wells. At the end of the assay, a graph and data sheet were generated in the system, allowing for the analysis of the effect of the different compounds on the mitochondrial function.

#### Dual lectin and immunocytochemistry

Lectin cytochemistry was carried out in combination with ICC to assess the glycosylation profile of the cells in each condition. Briefly, cells were fixed in 4% PFA (Sigma, Ireland) in PBS for 20 minutes and washed in Tris-buffered saline (TBS) supplemented with 1 mM of Mg2+ and Ca2+ (which are needed for lectins to bind) and in 0.05% triton X in TBS (TBS-T). Samples were blocked with a solution of 2% periodate-treated BSA (pBSA) in TBS for one hour at room temperature to prevent unspecific binding (23,48). The cells were then washed with TBS and incubated with FITC-conjugated SNA-I, MAA, DSL or GNA lectins (all from Vector Labs, UK) in the dark for one hour at room temperature. Inhibitory controls were also carried out at the same time in the presence of the respective haptenic sugar. Subsequently, the cells were washed in TBS and blocked again in 2% pBSA for one hour at room temperature. Afterwards the samples were incubated with primary antibodies (anti-β-tubulin III produced in rabbit (Sigma, Ireland, T2200, 1:400) and anti-GFAP produced in mouse (Sigma, Ireland, G3893, 1:400)) overnight, at 4 °C. On the following day, samples were washed in TBS-T and incubated with secondary antibodies at room temperature for one hour (AlexaFluor 594 goat anti-rabbit IgG (H+L) (Invitrogen, Ireland, A-11012, 1:500), Cy®5 goat anti-mouse IgG (H+L) (Invitrogen, A10524, 1:500)) in TBS. Finally, samples were washed with TBS-T and the nuclei stained with Hoechst (ThermoFisher, Ireland, 33342, 1:1000). Specimens were mounted on microscope cover slides with fluoromount aqueous mounting medium (Sigma, Ireland, F4680) and images were taken on PerkinElmer Operetta high content imaging system. The quantification of lectin expression was performed in the Operetta software, where the total area stained by the lectin in each image and respective intensity was calculated, and this was normalised by the total number of cells present on that same image.

#### RNA isolation and gene expression

Cells were washed thoroughly with PBS before adding TRIzol®. A pipette tip was used to scrape them and then the TRIzol® solution was transferred to a 2 mL microcentrifuge Eppendorf tube. Samples were centrifuged at 1,200 ×g for 5 min at 4 °C and the cellular debris was discarded. The clear supernatant was transferred to a clean tube and 200 μL chloroform was added per 1 mL TRIzol® solution, following the protocol described before for RNA extraction from tissue. cDNA synthesis was also carried out as above-mentioned. cDNA products were amplified using Fast SYBR® Green Master Mix (Applied Biosystems®, 4385612) and following specific primers (Sigma-Aldrich) (Table S6).Reactions were conducted in triplicate using StepOnePlus™ Real-Time PCR System (Applied Biosystems®. The results were analysed using the 2-ΔΔCt method and presented as fold change (relative gene expression) normalized to 18S and basal control.

### Statistics

Data were processed using GraphPad Prism8 software and reported as mean ± standard deviation, if not otherwise stated. Comparisons among groups were performed by one-way or two-way ANOVA, followed by Tukey’s multiple comparison post-hoc test. One-way (or two-way) ANOVA were employed after confirming that the distribution of the sample mean was normal (Shapiro-Wilk normality test) and the variances of the population of the samples were equal to one another (unpaired t test, F-test for homogeneity of variance).

## Study approval

All human samples were acquired from and associated clinical and neuropathological data were supplied by the Parkinson’s UK Brain Bank, funded by Parkinson’s UK, a charity registered in England and Wales (258197) and in Scotland (SC037554). All of these were post-mortem samples, collected according to the ethical legislation in place in the UK. All experiments were done in accordance with the NUI Galway Research Ethic Committee guidelines, under the reference of 18-Mar-20.

