## Supplementary material for "Changes in Protein *N*-Glycosylation Regulation Occur in the Human Parkinsonian Brain in a Region-Specific Manner": Suppp file

**This file includes:**

Supporting text
Supporting materials and methods
Figures S1 to S2.
Tables S1 to S6.
Legend for Dataset described in Tables S7 and S8.
SI References

**Supporting information text:**

***N*-Glycome profile of the healthy human nigro-striatal regions is mainly composed of galactosylated, highly fucosylated and oligomannosylated structures**

After looking at the overall glycosylation traits, a more in-depth study of the *N*-glycosylation patterns was performed (list of the samples used is described in Table S2; healthy controls were 1:2 male to female ratio, and the ages were 79 ± 10 years). For this, a combination of hydrophilic interaction liquid chromatography – ultra performance liquid chromatography (HILIC-UPLC) and liquid chromatography-mass spectrometry (LC-MS) was used after assessing the reproducibility of the results with control samples (Figure S1). The combination of both approaches enabled the) identification and quantification of both low- and high-abundance of over 120 *N*-glycan isomers for both striatum and substantia nigra spread over 59 glycan peaks (GP) (Table S7, Table S8) (main glycan structures seen in each GP are described in Figure S2).

To analyse the distinct traits, the major glycan constituent of each peak was considered the representative feature of that peak and the main traits were calculated by adding the abundance of the peaks where the main glycan expressed shared the same structural feature (Table S1). The glycoproteins in the healthy striatum and substantia nigra were seen to carry a high percentage of galactosylated and core fucosylated structures (around 47% each). Additionally, approximately 81% of the total *N*-glycans are neutral in the striatum, whereas in the substantia nigra this value drops to around 73%. From the sialylated glycans, only about 2% and 5% contain polysialic acid in the striatum and substantia nigra, respectively (Figure 3). The abundance of bisected glycans is lower (around 14% in the striatum and 11% in the substantia nigra). Most of the glycans are also branched (approximately 68% in both regions) indicating a high complexity in the structures present in the nigrostriatal pathway (Figure 3).

Interestingly, alpha-galactose epitopes (Gal-α1-3-Gal) were detected (both by HILIC-UPLC and lectin array). However, since humans cannot synthesise this residue as they lack the enzyme α1,3galactosyltransferase, this is most probably of non-human origin, being possibly contamination or due to dietary residues.

**Supporting Materials and Methods**

**Dual lectin and immunohistochemistry**

Frozen tissue sections (10 μm thick) were warmed at room temperature for at least 45 minutes before beginning the staining. Sections were washed with 0.05% Triton-X (TBS-T) in Tris-buffered saline supplemented with Ca2+ and Mg2+ (TBS: 20 mM Tris-HCl, 100 mM NaCl, 1 mM CaCl2, 1 mM MgCl2, pH 7.2) three times during three minutes with gentle shaking. Then these sections were blocked with 3% periodate-treated high grade bovine serum albumin (pBSA) made in TBS for one hour at room temperature to block unspecific binding. The sections were then washed again three times with TBS for three minutes each wash. Afterwards, they were incubated with fluorescently labelled lectins (Table S4) prepared in TBS-T for one hour in the dark at room temperature. After washing the sections again with TBS, these were blocked again with 3% pBSA for one hour at room temperature and then incubated with primary antibodies (anti-glial fibrillary acid protein (GFAP) produced in rabbit (Dako, USA, Z0334, 1:400) or anti-Ionized calcium binding adaptor molecule 1 (Iba1) produced in rabbit (Wako, USA, 019-1941, 1:500)) overnight at 4oC. After 24h, sections were washed in TBS-T and TBS and incubated with the secondary antibody at room temperature for one hour (AlexaFluor 594 goat anti-rabbit IgG (H+L) (Invitrogen, Ireland, A-11012, 1:1000) in TBS. These were then washed with TBS-T and TBS and finally counterstained with Hoechst (ThermoFisher, Ireland, 33342, 1:2000 in TBS) for 15 minutes. Subsequently the sections were washed again in TBS and incubated with 0.2% sudan black B solution (since this is effective in quenching the autofluorescence from lipofuscin, a substance consisting of oxidised lipids and common in aged brains) in 70% ethanol to quench lipofuscin auto fluorescence (granules that accumulate with age in the human brain). After washing twice with TBS, sections were mounted using fluoromount aqueous mounting medium (Sigma, Ireland, F4680) and coverslips and imaged within three days after curing. Inhibition by the appropriate haptenic sugar was used as control for all lectins in the study.

**Lectin microarray**

Lectin solutions and antibody solutions were prepared in buffer (1.0mM D-glucose in PBS containing 0.01% of cy3-conjugated BSA). These dilutions was printed onto NHS functionalised glass slide. The slides containing the lectin arrays were incubated. The remaining NHS groups were quenched by placing the slides in a 30mM ethanolamine solution in borate buffer and then blocked with a 0.3 mg/mL BSA, 0.3mM $Ca2+$ solution in PBS-T 0.05%. The slides were directly dried by centrifugation without previously washing. All the lectins used in the microarray are listed in Table S3. To prevent possible interference from the sample buffer (RIPA® buffer (R0278, Sigma, Ireland) and cOmpleteTM Protease Inhibitor Cocktail (5056489001, Roche, Ireland)), during the labelling step the buffer was changed using 0.5mL 10kDa Amicon™ spin filters (Merck, Ireland). 30 µg of each sample was added to the spin filter and washed twice with PBS by centrifugation (Beckman Coulter Allegra X-22R Centrifuge). Glycoproteins were recovered at a final concentration of 0.3 µg/µL in PBS. A volume of 10 µL of PBS 10x was added to the 0.3 µg/µL solution of glycoprotein in PBS prepared previously (100µL). Afterwards, the solution was incubated with Alexa-555-NHS for 1h at room temperature. The excess dye was quenched by the addition of Tris buffer and the fluorescently labelled glycoprotein solutions were directly used in the lectin array analysis. The glycoprotein samples (at a concentration of 3 µg/mL) were added to the corresponding wells (previously printed on the slides) and incubated for 1h 30 at room temperature. The sample concentration was adjusted by adding lectin incubation buffer to the labelled glycoprotein solutions after labelling. Labelled-BSA and control dye samples were incubated under the same conditions as negative controls for lectin binding. Then the glycoprotein solutions were removed and the slide washed with PBS, dried by centrifugation and scanned on a G265BA microarray scanner (Agilent Technologies). The images obtained were analysed with Pro Scan Array Express® software (PerkinElmer). Two-way ANOVA was performed, followed by Tukey's post-hoc test and statistical significance set at *p<0.05 and **p<0.01.

***N*-glycan analysis**

Exoglycosidase digestions of 2-AB labelled *N*-linked glycans

The 2-AB labelled *N*-glycans were digested in a volume of 10 µL for 18h at 37oC in 50 mM sodium acetate buffer, pH5.5 (except jack bean α-mannosidase (JBM), which was in 100 mM sodium acetate, 2 mM Zn2+, pH 5.0). The enzymes used were: Arthrobacter urefaciens sialidase (ABS), 0.5U/mL; Streptococcus pneumoniae sialidase (NAN1), 5U/mL; bovine testes β-galactosidase (BTG), 1U/mL; bovine kidney α -fucosidase (BKF), 1U/mL; almond meal α-fucosidase (AMF),

400U/mL; β-N-acetylglucosaminidase cloned from Streptococcus pneumonia (GUH), 400U/mL; JBM, 400U/mL; Streptococcus pneumoniae β-galactosidase (SPG), 0.4U/mL; coffee bean α-galactosidase (CBG), 25U/mL. Enzymes were purchased from Prozyme (San Leandro, CA, USA) (ABS, NAN1, BTG, SPG, CBG, BKF, JBH) or New England Biolabs (Hitchin, Herts, UK) (AMF, JBM, GUH). After incubation, enzymes were removed by filtration through 10kDa MWCO microcentrifuge filtration tubes (Pall Corporation, NY, USA). Digested N-glycans from the different samples were analysed by HILIC-UPLC(3).

Liquid chromatography-mass spectrometry (LC-MS) of *N*-linked glycans

For liquid chromatography fluorescence quadrupole time-of-flight mass spectrometry (UPLC-FLR-QTOF MS) analysis, the dried N-glycan samples were reconstituted in 3 µL of milliQ water and 9 µL acetonitrile. Online coupled fluorescence (FLR)-mass spectrometry detection was performed using a Waters Xevo G2 QTof with AcquityTM UPLC (Waters Corporation, Milford, MA, USA) and BEH Glycan column (1.0 x 150mm, 1.7 µm particle size). Sample injection volume was 10 µL. The flow rate was 0.150 mL/min and column temperature was kept at 60°C. Solvent A was 50 mM ammonium formate (pH 4.4) and solvent B was acetonitrile. A 40 minute linear gradient was used and was as follows: Solvent A at 28% for one minute, increase from 28% to 43% for 30 minutes, increase from 43% to70% for one minute, constant at 70 % for three minutes, decrease from 70% to 28% for one minute and constant at 28% for four minutes. To avoid contamination of the system, the flow was sent to waste for the first 1.2 minutes and after 32 minutes. The fluorescence detector settings were as follows: λexcitation: 320 nm, λemission: 420 nm; data rate was 1 pts/second and a PMT gain = 10. For mass spectrometry (MS) acquisition data, the instrument was operated in negative-sensitivity mode with a capillary voltage of 1.8 kV. The ion source block temperature was set at 120oC and the nitrogen desolvation gas temperature was set at 400°C. The desolvation gas had a flow rate of 600 L/h. The cone voltage was kept at 50 V. Full-scan data for glycans were acquired over m/z range of 450 to 2500. Data was collected and processed using MassLynxTM 4.1 software (Waters Corporation, Milford, MA, USA).

Glycan nomenclature

All *N*-glycans share the same pentasaccharide core composed of two core GlcNAcs linked to three mannoses. F at the start of the abbreviation indicates a core α(1,6)-fucose linked to the inner GlcNAc. Otherwise, F indicates an outer arm α(1,3) or α(1,4)-fucose linked to antenna or galactose. Mx indicates the number (x) of mannose residues on the core GlcNAcs. Ax refers to the number (x) of GlcNAc (antenna) on the trimannosyl core. Gx relates to the number (x) of β(1,4)-linked galactose on the antenna and Galx to the number (x) of α(1,3/4/6)-linked galactose on β(1,4)-linked galactose. Sx stands for the number (x) of sialic acids (neuraminic acids) linked to galactose, through an α(2,3)-, α(2,6)- or α(2,8)-linkage depending on the number inside the parentheses (3, 6 or 8, respectively). Sgx concerns the number (x) of glycolylneuraminic acids linked to galactose, and the number in parentheses corresponds to the linkage as before. Lacx indicates the number (x) of poly-N-Acetyllactosamine repeats containing GlcNAc linked β(1,4)- to galactose.

**DMB assay**

To assess the type of sialic acids present in the brain glycoproteins, a kit was used to release the sialic acids and label them with 1,2-diamino-4,5-methylenedioxybenzene Dihydrochloride (DMB). After homogenising the tissue as described previously, the glycoprotein solution was dried in a SpeedVac (Savant™ SPD131DDA SpeedVac™ Concentrator, ThermoFisher) overnight. On the next morning, 25 µL of 2M acetic acid solution was added to each sample, to the controls and the standards. These were briefly vortexed and centrifuged. Then they were incubated at 80oC for two hours and cooled to room temperature afterwards. 5 µL from each vial was kept at -20oC for the following steps.For the sample labelling, 440µL of the mercaptoethanol solution was added to the vial of sodium dithionite and mixed until dissolved. This solution was then added to the DMB dye and mixed. Afterwards, 20µL of this labelling reagent was added to each sample, controls and standards, mixed thoroughly and briefly centrifuged. Then the vials were incubated for three hours at 50oC in the dark. The reaction was terminated by the addition of 475 µL of water to each sample and control or of 480 µL of water to each sialic acid standard.

Ultra-high performance liquid chromatography (UHPLC)

UHPLC was performed to analyse these samples using the LudgerSepTM uR2 UHPLC column, 175 A, 1.9 µm silica derivatised particles with octadecylsiliane coating, 2.1 x 100mm (Luger, UK) on an Acquity system. Solvent A was acetonitrile:methanol:water (9:7:84) and solvent B was acetonitrile. The sample temperature was set to 10oC and the column temperature to 30oC. The method had a 15 minute duration, the flow rate was constant at 0.25 mL/min. During the first 7 minutes, the flow was solely composed of solvent A. This was reduced to 10 % (and solvent B increased to 90%) between 7.5 and 8 minutes, and returned to the initial values between minute 8.5 and 15. N-acetylneuraminic acid quantitative standard, N-glycolylneuraminic acid quantitative standard and a sialic acid reference panel containing Neu5Ac, Neu5Gc, Neu5,7Ac2, Neu5Gc,9Ac, Neu5,8Ac2, Neu5,9Ac2 and Neu5,x,xAc3 (where x is an unknown acetyl position) were used at the beginning of the run to calibrate the system. All of these were included in the kit. A negative control with only buffer (without sample) was also used.

**Matrix-assisted laser desorption/ionisation mass spectrometry imaging (MALDI-MSI) of *N*-glycans**

For spatial *N*-glycome analysis through MALDI mass spectrometry imaging, frozen 10 µm brain tissue sections on SuperFrostTM Plus Adhesion charged slides (Fischer Scientific, Ireland) were thawed and dehydrated in serial dilutions of ethanol (70%, 90%, 100%, 100%) for two minutes each. Then, sections were incubated at 60 oC for 50 minutes, followed by delipidation in Carnoy's solution twice (30% chloroform, 10% glacial acetic acid, 60% ethanol), for three minutes, and a wash in running tap water. Finally, these were incubated with 10% formalin solution neutral buffered (Sigma, Ireland) (30 minutes), followed by two washes in tap water. The samples were then air-dried and kept in a desiccator until further analysis. The transformed slides were exposed to antigen retrieval using citraconic anhydride buffer (pH3), prepared by mixing 25 µL of citraconic anhydride (Sigma, Germany) in 50 mL of HPLC grade water. Slides were incubated in this buffer in a vegetable steamer (approximately 95 oC) for 30 minutes and then washed in serial dilutions of the buffer by replacing half of the buffer with HPLC grade water (three times), eventually switching it completely with water. The slides were air-dried and scanned before applying PNGase F. After antigen retrieval, slides were coated with an aqueous solution of recombinant PNGaseF (Serva, Germany) at 0.1 µg/ µL at 45 oC, using an HTX TM-SprayerTM (HTX Imaging, USA) as previously described (4). This was followed by an incubation of two hours at 37 oC in a humidified chamber and placed in the desiccator until sprayed with matrix on the same day. α-Cyano-4-hydroxycinnamic acid (CHCA) matrix was prepared fresh (7 mg/mL in 50% acetonitrile 0.1% TFA) and applied in the sections at 100 µL/ min at 79 oC using an HTX TM-SprayerTM. Coated slides were stored in a desiccator until being analysed (no longer than 48 h post-spraying).

**Western blot**

Protein lysates from brain samples previously homogenised in RIPA buffer (R0278, Sigma, Ireland) and cOmpleteTM Protease Inhibitor Cocktail (5056489001, Roche, Ireland, 1:25) were boiled at 95oC for seven minutes with loading buffer containing 1% bromophenol blue and 200mM of DTT. Equal amounts (5 µg or 10 µg, depending on the detection antibody) of protein samples were run on an SDS polyacrylamide gel with 12% or 15% acrylamide depending on the size of the proteins to be detected. After the run, the proteins were transferred onto a 0.45 NC nitrocellulose membrane (Fisher Scientific, Ireland) using the Trans-Blot® TurboTM transfer system (Biorad, UK). Ponceau red S staining was performed to confirm the presence of the proteins in the membrane. The ponceau red was washed off with 5% acetic acid and then the membrane was washed with TBS-T (tris buffered solution with 0.05% Tween20). The membrane was blocked with 5% milk in TBS-T for one hour at room temperature. This was followed by incubation with primary antibody for GRP94 (Cell Signalling Technology, The Netherlands, 2104S, 1:1000), PDI (Cell Signalling Technology, The Netherlands, 3501S, 1:1000), GRP78 (Abcam, UK, ab32618, 1:3000), ATF6 (Abcam, UK, ab37149, 1:750), phospho-eIF2α (Cell Signalling Technology, The Netherlands, 9721S, 1:1000), ATF4 (Cell Signalling Technology, The Netherlands, 11815S, 1:1000) and CHOP (Cell Signalling Technology, The Netherlands, 2895S, 1:1000) overnight at 4oC. On the next day, the membrane was washed three times with TBS-T and incubated with the appropriate horseradish peroxidase-conjugated secondary antibody (Goat anti-mouse IgG, Fisher Scientific, Ireland, 31430, 1:10000, or goat anti-rabbit IgG, Fisher Scientific, Ireland, 31460, 1:10000) for one hour at room temperature. After three washes in TBS-T, signals were detected using SuperSignalTM WestPico PLUS ECL (Fisher Scientific, Ireland, 34577). Quantification of the signal intensity was performed using Image Studio Lite and normalised by the expression of β-actin (which was carried out in all the blots performed). If the data was normally distributed, one-way ANOVA was performed, followed by Dunnett post-hoc test, and statistical significance was set at *$p<0.05$. If the data was not normally distributed, a Kruskall-Wallis test followed by a Dunn's post hoc test were carried out, and statistical significance was set at *$p<0.05$.

**Pathological analysis**

Immunohistochemistry (chromogenic staining)

Frozen tissue sections (10 µm thick) were warmed at room temperature for at least 45 minutes before starting the staining. Sections were incubated with 0.3% hydrogen peroxidase in 70% MeOH in PBS for 20 minutes to block endogenous peroxidase. Then the sections were washed twice with PBS and blocked with 5% normal goat serum (NGS) in PBS-T (PBS with 0.2% Triton X) for one hour at room temperature. The slides were then incubated with primary antibody solution prepared in 5% NGS in PBS-T (anti-GFAP produced in rabbit (Dako, USA, Z0334, 1:400) or anti-Iba1 produced in rabbit (Wako, USA, 019-1941, 1:500)) overnight at 4 oC. On the next day the sections were washed twice with PBS and incubated with the biotinylated secondary antibody prepared in PBS for 30 minutes at room temperature. After washing twice in PBS, slides were incubated with streptavidin ABC HRPcomplex (VectorLabs®, UK, PK-6100) for 30 minutes at room temperature. Then the slides were washed and incubated with 3,3'-Diaminobenzidine tetrahydrochloride hydrate (DAB) (2 mg/mL) (Sigma, Ireland, D9015) activated with 30% hydrogen peroxide for five minutes. Finally, the sections were counterstained with Heamatoxylin Gill no.2 (Sigma, Ireland, GHS216), dehydrated in ethanol, cleared in xylene and mounted with DPX mountant (Sigma, Ireland, 06522).

Haematoxylin/Eosin

Frozen tissue sections (10 μm thick) were warmed at room temperature for at least 45 minutes before the start of staining. Sections were hydrated in serial dilutions of ethanol (100%, 95%, 90%, 70%, 50% EtOH), followed by tap water. Then the sections were incubated with Heamatoxylin Gill no.2 (Sigma, Ireland, GHS216) followed by Eosin Y (Sigma, Ireland, HT110132). Finally, the slides were dehydrated in three solutions of 100% EtOH, cleared in xylene and mounted with DPX® mountant (Sigma, Ireland, 06522).

Terminal deoxynuclotidyl transferase dUTP nick end labelling (TUNEL) assay

TUNEL assay was performed using the Kit HRP-DAB from Abcam (Abcam, UK, ab206386) and the manufacturer's instructions were followed. Briefly, the frozen sections were warmed at room temperature for at least 45 minutes, permeabilised with Proteinase K (1:100) for ten minutes and washed with TBS. Sections were incubated with 3% hydrogen peroxidase in MeOH for five minutes to block endogenous peroxidase, washed with TBS, equilibrated with TdT equilibration buffer for 30 minutes and incubated with TdT labelling reaction mix in a humidified chamber at 22oC for 1h 30. The reaction was stopped with the stop buffer and the slides were washed with TBS. The sections were then blocked with blocking buffer for ten minutes at room temperature and incubated with the conjugate solution for 30 minutes at room temperature. Slides were washed with TBS and the stain developed using DAB for 15 minutes. Finally, the sections were counterstained with methyl green, dehydrated in two solutions of 100% EtOH, cleared in xylene and mounted with DPX® mountant (Sigma, Ireland, 06522).

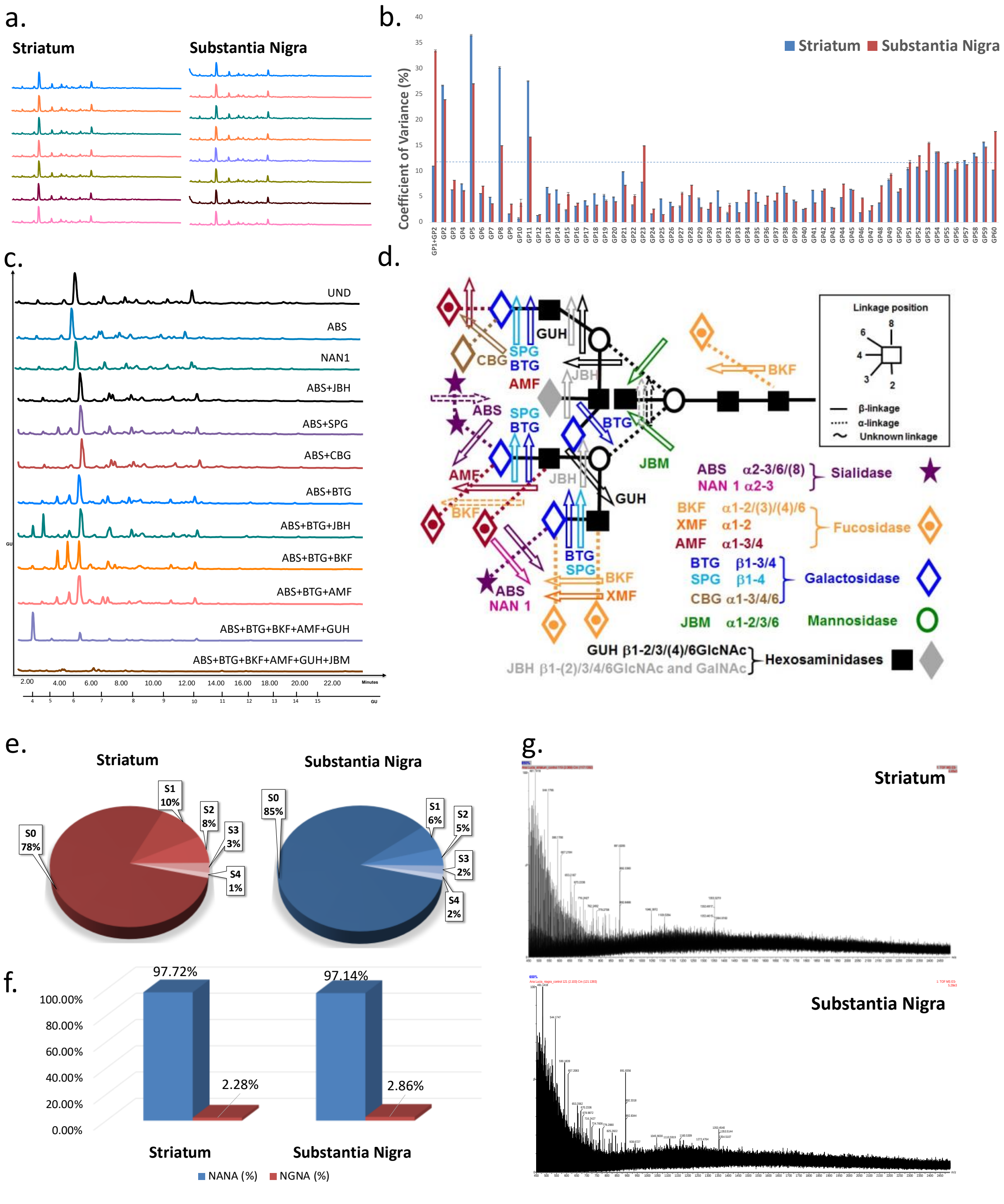


**Figure S1. Development and optimisation of the glyco-analytical platform based on liquid-chromatography methods to characterise the human nigro-striatal *N*-glycome**. **a.** Reproducibility of the platform is represented by the overlay of biological replicates, which are indicated by different colours for both healthy striatum and substantia nigra. **b.** Comparison of the individual glycan peaks (GP) area coefficients of variance for eight samples from both striatum and substantia nigra using HILIC-UPLC. Separation of chromatograms into GP can be seen in Figure **S2**. To characterise the chromatograms obtained in each region, a sequence of exoglycosidase digestions was performed **c.** Representative chromatograms for the exoglycosidase digestions performed on the striatum samples using HILIC-UPLC (similar profiles were obtained for substantia nigra – data not shown). Through the shift of each peak after incubation with the different enzymatic cocktails, it was possible to build the glycan structures present in each of the peaks, as well as to calculate their abundance (Figure **S2**). Detailed information on all exoglycosidase digestions can be found in **Supplementary table S7**. **d.** Schematic representation of the specificity of each exoglycosidase. **e.** Percentage fractions of sialylated *N*-glycans in both striatum and substantia nigra, classified according to the degree of sialylation (mono- (S1), di-(S2), tri- (3), tetra- (S4)) using WAX-UPLC. **f.** Percentage fraction of the different types of sialylation (Neu5Ac (NANA) vs Neu5Gc (NGNA)) present in the overall pool of glycoproteins in the healthy striatum and substantia nigra using DMB assay. Human cells no longer have the ability to synthesize Neu5Gc, however it is possible that these were included in the glycan structures due to dietary incorporation (1). **g**. LC-MS profiles of the *N*-glycans from both healthy striatum and substantia nigra. All the results gathered from the exoglycosidase digestions, WAX-UPLC profile, DMB profile and LC-MS were used to characterise and describe the *N*-glycome HILIC-UPLC-based profile for each brain region, in healthy conditions and upon disease. Abbreviations: DMB: 1,2-diamino-4,5-methylenedioxybenzene Dihydrochloride, LC-MS: Liquid chromatography-mass spectrometry, HILIC-UPLC: Hydrophilic Interaction Liquid Chromatography - Ultra Performance Liquid Chromatography, WAX-UPLC: Weak Anion Exchange - Ultra Performance Liquid Chromatography.

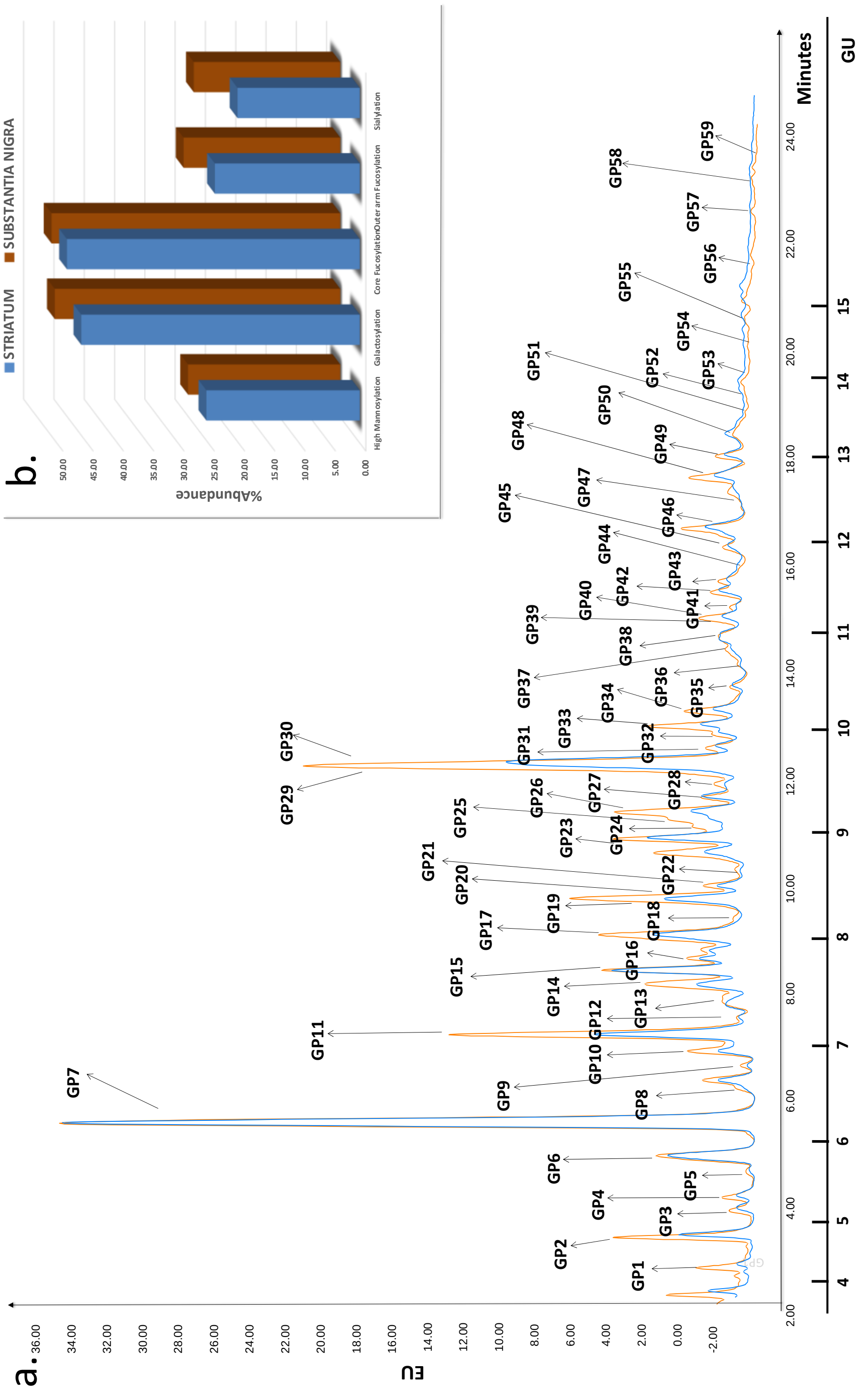
a.
EU
Minutes
GU
b.
STRIATUM
SUBSTANTIA NIGRA
%Abundance
High Mannosylation
Galactosylation
Core Fucosylation
Outer arm Fucosylation
Sialylation

c.

| Peak number | STRIATUM | | | | Peak number | SUBSTANTIA NIGRA | | | |
|---|---|---|---|---|---|---|---|---|---|
| | HEALTHY | | PD | | | HEALTHY | | PD | |
| GP | Structure | Peak Area (%) | Structure | Peak Area (%) | GP | Structure | Peak Area (%) | Structure | Peak Area (%) |
| 1 | M3 | 0.44 | M3 | 0.37 | 1 | M3 | 0.49 | M3 | 0.43 |
| 2 | F(6)M3 | 2.11 | F(6)M3 | 1.53 | 2 | F(6)M3 | 1.70 | F(6)M3 | 1.76 |
| 3 | M4 | 0.43 | M4 | 0.46 | 3 | M4 | 0.42 | M4 | 0.44 |
| 4 | F(6)A1 | 0.48 | F(6)A1 | 0.53 | 4 | F(6)A1 | 0.51 | F(6)A1 | 0.59 |
| 5 | A2B | 0.78 | A2B | 0.20 | 5 | M4A1 | 0.32 | M4A1 | 0.31 |
| 6 | A3 | 2.57 | A3 | 2.53 | 6 | F(6)A2 | 2.29 | A3/F(6)A2 | 2.27 |
| 7 | F(6)A3 (M5) | 20.46 | F(6)A3 (M5) | 21.09 | 7 | F(6)A3 (M5) | 19.01 | F(6)A3 (M5) | 18.74 |
| 8 | F(6)M5 | 1.21 | F(6)M4A1B | 1.19 | 8 | F(6)M4A1B | 1.16 | F(6)M5 | 1.24 |
| 9 | M5A1 | 0.15 | A4B | 0.16 | 9 | A4B | 0.15 | A2G1GalNAc1S(3)1 (SO4- on GalNAc) | 0.17 |
| 10 | F(6)A4 | 1.05 | F(6)A4 | 1.16 | 10 | F(6)A4 | 1.08 | F(6)A4 | 1.28 |
| 11 | M6 | 5.09 | M6 | 5.13 | 11 | M6 | 5.81 | M6 | 6.18 |
| 12 | M5A2 | 0.40 | M5A2 | 0.43 | 12 | M5A2 | 0.35 | A2F1G1GalNAc1 | 0.33 |
| 13 | A3G(4)2/A3G(3)2 | 0.81 | F(6)A2F1G(4)1 | 0.82 | 13 | F(6)A2BG1 | 0.68 | F(6)A4GalNAc1 | 0.62 |
| 14 | M7/A4F1 | 1.79 | M7 | 1.84 | 14 | A3BG(4)2 | 2.02 | F(6)A2G(3)2/F(6)A2G(4)2 | 2.07 |
| 15 | F(6)A2BF1G1 | 3.89 | F(6)A3F1G1/F(6)A2BF1G1 | 3.98 | 15 | F(6)M5A2 | 3.37 | F(6)A3F1G1/F(6)A2BF1G1 | 3.09 |
| 16 | M7 | 1.46 | F(6)A3F1G1 | 1.55 | 16 | M7 | 1.36 | M7/F(6)A2F1G1GalNAc1 | 1.35 |
| 17 | M7 | 4.42 | M7 | 4.42 | 17 | M7 | 4.68 | M7 | 4.72 |
| 18 | F(6)A4BF1 | 0.95 | F(6)A4GalNAc2 | 1.03 | 18 | F(6)A2BG1S(3)1 | 0.92 | M7 | 0.91 |
| 19 | A3F3 | 1.18 | A3F3 | 1.41 | 19 | F(6)A4F1GalNAc1 | 1.29 | F(6)A4F1GalNAc1 | 1.56 |
| 20 | A3G(4)3 | 1.09 | A2G2S(3)1 | 0.90 | 20 | A2G2S(3)1 | 1.27 | A2G2S(3)1 | 1.09 |
| 21 | F(6)A4G1Lac1 | 1.12 | F(6)A3BF1G(4)1 | 1.12 | 21 | F(6)M7 | 1.02 | A3F1G1Sg(6)1 | 0.98 |
| 22 | A3BG(3)3/A3BG(4)3 | 0.50 | A3BG(3)3/A3BG(3)3 | 0.52 | 22 | A3BG(3)3 | 0.48 | F(6)A2G(4)2S(3)1 | 0.51 |
| 23 | M8 | 4.15 | M8 | 4.30 | 23 | M8 | 4.50 | M8 | 4.35 |
| 24 | M8 | 0.46 | M8 | 0.51 | 24 | M8 | 0.44 | F(6)A4F2G(3)4 | 0.35 |
| 25 | F(6)A3BG(4)3 | 0.95 | F(6)A4F1G2 | 0.98 | 25 | F(6)A3BG(4)3/A4F2G1 | 1.00 | M8 | 1.02 |
| 26 | F(6)A2BF2G2 | 1.85 | F(6)A2BF2G2 | 1.90 | 26 | F(6)A3F2G1S(3)1/F(6)A3F2G1S(6)1 | 2.02 | F(6)A3F2G1S(3)1 | 1.89 |
| 27 | A3BF2G1S(3)2/A3BF2G1S(6)2 | 1.86 | F(6)A4F3 | 1.95 | 27 | F(6)A4F3 | 1.72 | A3BF2G1S(6)2 | 1.44 |
| 28 | A4G(3)3Lac1/A4G(4)3Lac1 | 0.93 | F(6)A2F2G2GalNAc1 | 0.89 | 28 | F(6)A3G2Gal2 | 0.88 | A4G(3)3Lac1/A4G(4)3Lac1 | 0.81 |
| 29 | M9 | 5.05 | M9 | 5.43 | 29 | M9 | 5.31 | M9 | 4.84 |
| 30 | A2G2S(6)2 | 3.17 | A2G2S(3)2 | 3.13 | 30 | A2G2S(6)2 | 3.66 | F(6)A2G(4)2S(3,6)2 | 3.87 |
| 31 | F(6)A4G3Lac1 | 0.98 | A3F1G1Gal2GalNAc1 | 1.05 | 31 | A4G(4)4/A4G(3)4 | 1.06 | A4G(4)4 | 1.21 |
| 32 | F(6)A3G3S(3)1 | 0.64 | N/A | 0.61 | 32 | FA4F1GalNAc1S(6,8)2 | 0.66 | F(6)A3G3S(3)1 | 0.55 |
| 33 | F(6)A2F2G2S(6)1 | 1.77 | A4BG3S(3)1 | 1.81 | 33 | A4F2G2 | 1.71 | A4BG(3)4 | 1.57 |
| 34 | F(6)A4F3G1 | 1.08 | F6)A3G2Lac2 | 0.94 | 34 | F(6)A4F3G(4)1 | 0.99 | F(6)A4F3G1 | 0.96 |
| 35 | F(6)A3BG2Lac2 | 0.78 | F(6)A4G(4)4 | 0.85 | 35 | A3BG3S(6)2 | 0.81 | F(6)A4G2Sg(6)1 | 0.74 |
| 36 | F(6)A4F1G(3)3 | 0.43 | F(6)A4F1G(3)3 | 0.44 | 36 | F(6)A4F1G3 | 0.40 | F(6)A4F1G3 | 0.29 |
| 37 | A4G(3)4Gal1 | 0.52 | A4G3GlcNAc3 | 0.54 | 37 | A4G4Gal1 | 0.51 | A4G(4)4Gal1 | 0.55 |
| 38 | M10 | 2.20 | F(6)A3G3S(3)2 | 2.28 | 38 | F(6)A3G3S(6)2 | 1.97 | F(6)A3F2G(3)3/F(6)A3F2G(4)3 | 1.78 |
| 39 | F(6)A3BG3GlcNAc2S(3,6)2/ F(6)A3BG3GlcNAc2S(6)2 | 0.78 | F(6)A4F3GalNAc3 | 0.65 | 39 | A4BF3G1S(6)1 | 1.05 | A4BF3G1S(6)1 | 1.07 |
| 40 | F(6)A4BF4G1 | 0.71 | F(6)A4BF4G1 | 0.79 | 40 | F(6)A4BF4G1 | 0.81 | F(6)A4BF4G1 | 0.93 |
| 41 | F(6)A4G4Gal1 | 0.47 | F(6)A4G(4)4Gal1 | 0.45 | 41 | F(6)A4G(4)4Gal1/F(6)A4G(3)4Gal1 | 0.57 | F(6)A4F3GalNAc4 | 0.63 |
| 42 | M11 | 0.97 | A3F1G1Gal2S(3)2/A3F1G1Gal2S(6)2 | 1.02 | 42 | M11 | 1.04 | F(6)A4G2Sg(3)2 | 1.01 |
| 43 | A3F1G1Gal2S(3)3 | 1.00 | F(6)A4F1G2S(6)2 | 1.00 | 43 | A3F1G1Gal2S(3)3 | 1.00 | A3F1G1Gal2S(3)3 | 1.11 |
| 44 | F(6)A3F1G2Sg(3,3,8)3 | 0.35 | F(6)A3F1G2Sg(3,3,8)3 | 0.41 | 44 | A4F3G4 | 0.30 | A4F3G(4)4 | 0.22 |
| 45 | F(6)A4G(3)4Lac1Gal1/ F(6)A4G(4)4Lac1Gal1 | 1.38 | F(6)A4F3G1Sg(3)1 | 1.35 | 45 | F(6)A4G(3)4Lac1Gal1/ F(6)A4G(4)4Lac1Gal1 | 1.42 | F(6)A4G3Lac1Gal1 | 1.66 |
| 46 | F(6)A4F3G3 | 1.60 | A4BG3S(6)1Sg(6)1 | 1.51 | 46 | F(6)A3BF1G2S(3,6,8)3,F(6)A3BF1G2S(3,3,8)3 | 2.10 | F(6)A3BF1G2S(3,3,8)3/F(6)A3BF1G2S(6,6,8)3 | 1.49 |
| 47 | A4BF3G3 | 0.36 | A4BG3Lac1GlcNAc2S(3)1 | 0.32 | 47 | F(6)A3F2G1S(3,8,8,8)4 | 0.80 | F(6)A4F3G1S(6,8)2 | 1.22 |
| 48 | F(6)A4F3G4 | 2.20 | F(6)A4F(2)3G(4)4 | 2.17 | 48 | F(6)A2G2Gal2Sg(6)2 | 1.94 | F(6)A2G2Gal2Sg(6)2/F(6)A4F1G3S(3)3/ F(6)A4F1G3S(6)3 | 2.26 |
| 49 | A3F1G(4)3S(6)2Sg(3)1 | 1.18 | F(6)A3BG3Sg(6)2 | 1.08 | 49 | A3F(3)1G(4)3S(6)2Sg(3)1 | 1.81 | A3F(3)1G(4)3S(6)2Sg(3)1 | 1.54 |
| 50 | A4G4S(3)4 | 0.40 | A4G4S(3)4 | 0.39 | 50 | F(6)A4BF4G1Sg(6)1/A4BF3G1S(3,8,8)3 | 1.99 | A4G4S(3)4/F(6)A4BF4G1Sg(3)1 | 0.57 |
| 51 | F(6)A4G(3)4Lac3/F(6)A4G(4)4Lac3 | 2.15 | A4BF3G3GalNAc2 | 2.13 | 51 | A4F4G(4)4S(3)1 | 0.41 | A4F4G(4)4S(3)1 | 2.43 |
| 52 | F(6)A4G4S(3)4 | 0.81 | F(6)A4G4S(6)4 | 0.83 | 52 | F(6)A4G4S(3)4 /F(6)A4G4S(6)4 | 0.78 | A4G(3)4Gal1GalNAc2S(3)2/A4G(3)4Gal1GalNAc2S(6)2 | 0.65 |
| 53 | A4BG4S(3)3/A4BG4S(6)3 | 0.82 | A4BG4S(6)3 | 0.83 | 53 | A4BG4S(3)3/A4BG4S(6)3 | 0.87 | A4BG4S(3)3/A4BG4S(6)3 | 0.96 |
| 54 | A4F3G4S(3)4/A4F3G4S(6)4 | 1.35 | A4F3G4S(3)4/A4F3G4S(3,6,6,6)4/ A4F3G4S(6)4 | 1.33 | 54 | A4F3G4S(6)4 | 1.19 | A4F3G4S(6)4 | 1.04 |
| 55 | F(6)A4BF4G2Gal3 | 0.29 | F(6)A4BF4G2Gal3 | 0.25 | 55 | F(6)A4G4Lac4Gal1 | 0.25 | F(6)A4G4Lac4Gal1 | 0.43 |
| 56 | F(6)A4G4Lac3S(6)2 | 1.58 | F(6)A4G4Lac3S(3)2 | 1.55 | 56 | F(6)A4F3G4S(3,3,6)3/F(6)A4F3G4S(6)3 | 1.45 | F(6)A4G4Lac3S(3)2 | 1.33 |
| 57 | A2G2S(6,6,8,8)4 | 0.40 | A2G2S(6,6,8,8)4 | 0.41 | 57 | A2G2S(6,6,8,8)4 | 0.40 | A2G2S(6,6,8,8)4 | 0.40 |
| 58 | F(6)A4BF2G3S(6)3 | 0.45 | F(6)A4F3GalNAc1S(6,8,8,8)4 | 0.37 | 58 | F(6)A4BF2G(4)3S(6)3 | 0.42 | F(6)A4F2G4S(6)1Sg(3)4 | 1.11 |
| 59 | F(6)A4F2G4Sg(3)4S(8)1 | 0.57 | A4BF3G4S(6)4 | 0.54 | 59 | A4G4Gal1GalNAc2S(6)4 | 0.69 | A4G4Gal1GalNAc2S(6)4 | 0.06 |

d.

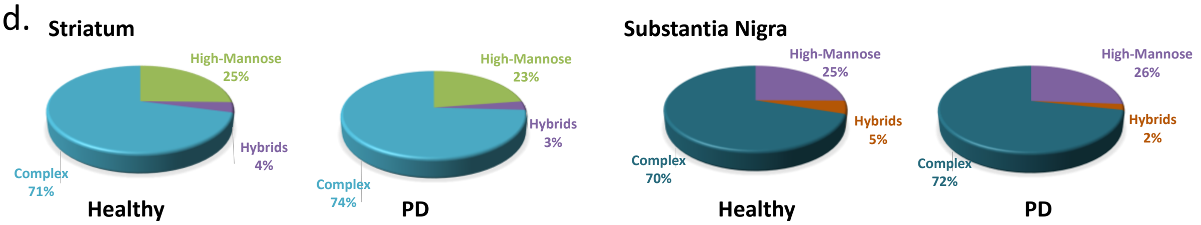


**Figure S2. *N*-glycome in nigro-striatal regions of the healthy controls and PD human patients.** a. HILIC-UPLC chromatograms obtained for both human striatum (blue profile) and substantia nigra (orange profile). The *N*-glycome for both these regions was separated into 59 chromatographic peaks (GP). EU – emission units. b. Abundance of the main glycosylation traits present in the healthy striatum and substantia nigra. c. Summary of the detailed composition analysis for each of these peaks, indicating the major glycan structure present in each peak and the peak area. d. Abundance of the three general types of *N*-glycans (hybrid, complex and high mannose) in both the striatum and substantia nigra, in healthy and PD samples

**Table S1. Calculation of derived glycosylation traits in the striatum.** The description of each glycosylation trait and the calculation rationale are presented as follows.

| | | | Healthy/ILBD | | PD | |
|---|---|---|---|---|---|---|
| | | | Striatum | Substantia Nigra | Striatum | Substantia Nigra |
| **Charge analysis** | N1 | The percentage of total neutral glycans in total glycans | SUM(GP1:17,GP19:26,GP28:29,GP31,GP34:38,GP40:42,GP45:48,GP51,GP55) | SUM(GP1:17,GP19,GP21:25,GP27:29,GP31,GP33:34,GP36:37,GP40:42,GP44:45,GP55) | SUM(GP1:19,GP21:29,GP31,GP34:37,GP39:41,GP48,GP51,GP55) | SUM(GP1:8,GP10:19,GP23:25,GP28:29,GP31,GP33:34,GP36:38,GP40:41,GP44:45,GP55) |
| | $S^{Total}$ | The percentage of total sialylated glycans in total glycans | SUM(GP18,GP27,GP30,GP32:33,GP39,GP43:44,GP49:50,GP52:54,GP56:59) | SUM(GP18,GP20,GP26,GP30,GP32,GP35,GP38,:39,GP43,GP46:54,GP56:59) | SUM(GP20,GP30,GP33,GP38,GP42:47,GP49:50,GP52:54,GP56:59) | SUM(GP9,GP20:22,GP26:27,GP30,GP32,GP35,GP39,GP42:43,GP46:54,GP56:59) |
| | S1 | The percentage of mono-sialylated glycans in total glycans | SUM(GP18,GP32:33) | SUM(GP18,GP20,GP26,GP39,GP49,GP50/2,GP51) | SUM(GP20,GP33,GP45,GP47) | SUM(GP9, GP20:22,GP26,GP32,GP39,GP50/2,GP51) |
| | S2 | The percentage of di-sialylated glycans in total glycans | SUM(GP27,GP30,GP39) | SUM(GP30,GP32,GP35,GP38,GP48) | SUM(GP30,GP38,GP42:43,GP46,GP49,GP56) | SUM(GP27,GP30,GP35,GP42,GP47,GP48/2,GP52,GP56) |
| | S3 | The percentage of tri-sialylated glycans in total glycans | SUM(GP43:44,GP49,GP53,GP56, GP58) | SUM(GP43,GP46,GP50/2,GP53,GP56,GP58) | SUM(GP44,GP53) | SUM(GP43,GP46,GP48/2,GP49,GP53) |
| | S4 | | SUM(GP50,GP52,GP54,GP57) | SUM(GP47,GP52,GP54,GP57,GP59) | SUM(GP50,GP52,GP54,GP57:59) | SUM(GP50/2,GP54,GP57,GP59) |
| | Phos | The percentage of phosphorylated glycans in total glycans | ND | ND | ND | ND |
| | $S^{Ac}$ | The percentage of acetylated charged species | ND | ND | ND | SUM(GP9) |
| | Total charged species | The percentage of total charged species in total glycans | SUM(GP18,GP27,GP30,GP32:33,GP39,GP43:44,GP49:50,GP52:54,GP56:59) | SUM(GP18,GP20,GP26,GP30,GP32,GP35,GP38,:39,GP43,GP46:54,GP56:59) | SUM(GP20,GP30,GP33,GP38,GP42:47,GP49:50,GP52:54,GP56:59) | SUM(GP9,GP20:22,GP26:27,GP30,GP32,GP35,GP39,GP42:43,GP46:54,GP56:59) |
| | Poly | The percentage of polysialic acids in total glycans | SUM(GP39,GP44,GP57,GP59) | SUM(GP32,GP46:47, GP50/2,GP57,GP59) | SUM(GP42,GP44,GP57:58) | SUM(GP43,GP46,GP47,GP57,GP58) |
| **Branching** | A1 | The percentage of monoantennary glycans in total glycans | SUM(GP4) | SUM(GP4) | SUM(GP4) | SUM(GP4) |
| | A1B | The percentage of bisecting monoantennary glycans in total glycans | ND | ND | ND | ND |
| | A2 | The percentage of biantennary glycans in total glycans | SUM(GP18,GP30,GP33,GP57) | SUM(GP6,GP20,GP30,GP48,GP57) | SUM(GP13,GP20,GP28,GP30,GP5 | SUM(GP6/2,GP9,GP12,GP14,GP16/2,GP20,GP22,GP30,GP48/2,GP57) |
| | A2B | The percentage of bisecting diantennary glycans in total glycans | SUM(GP5,GP15, GP26) | SUM(GP13,GP18) | SUM(GP5,GP15/2,GP26) | SUM(GP15/2) |
| | A3 | The percentage of triantennary glycans in total glycans | SUM(GP6, GP7/2,GP13,GP19:20,GP32,GP43:44,GP49) | SUM(GP7/2,GP26,GP28,GP38,GP43,GP47, GP49) | SUM(GP6,GP7/2,GP15/2,GP16, GP19,GP31,GP34,GP38,GP42, GP44) | SUM(GP6/2,GP7/2,GP15/2,GP21,GP26,GP32,GP38, GP43,GP49) |
| | A3B | The percentage of bisecting triantennary glycans in total glycans | SUM(GP22,GP25,GP27,GP35,GP39) | SUM(GP14,GP22,GP25/2,GP35,GP46) | SUM(GP21:22, GP49) | SUM(GP27,GP46) |

| | | | Healthy/ILBD | | PD | |
|---|---|---|---|---|---|---|
| | | | Striatum | Substantia Nigra | Striatum | Substantia Nigra |
| **Branching** | A4 | The percentage of tetraantennary glycans in total glycans | SUM(GP21,GP31,GP34,GP36:37,GP41,GP45:46,GP48,GP50:52,GP54,GP56,GP59) | SUM(GP10,GP19,GP25/2,GP27,GP31:34,GP36:37,GP41,GP44:45,GP51:52,GP54:56,GP59) | SUM(GP10,GP25,GP27,GP35:37,GP39,GP41,GP43,GP45,GP48,GP50,GP52,GP54,GP56,GP58) | SUM(GP10,GP13,GP19,GP24,GP28,GP31,GP34:37,GP41,GP44:45,GP47,GP48/2,GP50/2,GP51:52,GP54:56,GP58:59) |
| | A4B | The percentage of bisecting tetraantennary glycans in total glycans | SUM(GP40,GP47,GP53,GP55,GP58) | SUM(GP9,GP39,GP40,GP50/2,GP53,GP58) | SUM(GP9,GP33,GP40,GP46:47,GP51,GP53,GP55,GP59) | SUM(GP33,GP39:40,GP50/2,GP53) |
| **Bisects** | A1B, A2B, A3B, A4B | The percentage of bisecting glycans in total glycans | SUM(GP5,GP15,GP22,GP25:27,GP35,GP39:40,GP47,GP53,GP55,GP58) | SUM(GP8:9,GP13,GP14,GP22,GP25/2,GP35,GP39:40,GP46,GP50/2,GP53,GP58) | SUM(GP5,GP8:9,GP15/2,GP21:22,GP26,GP33,GP40,GP46,GP47,GP49,GP51,GP53,GP55,GP59) | SUM(GP15/2,GP27,GP46,GP33,GP39-40,GP50/2,GP53) |
| **Oligo-mannose** | Total oligo-mannose | The percentage of oligomannose structures in total glycans | SUM(GP1:3,GP7/4, GP8, GP11, GP14/2, GP16:17, GP23:24, GP29, GP38, GP42) | SUM(GP1:3,GP7/4,GP11,GP17,GP21,GP23:24,GP29,GP42) | SUM(1:3,GP8,GP11:12,GP14,GP17,GP23:24,GP29,GP7/4) | SUM(GP1:3,GP7/4,GP8,GP11,GP16/2,GP17:18,GP23,GP25,GP29) |
| | Lower order oligo-mannose | The percentage of oligomannose structures from M1-M5 in total glycans | SUM(GP1:3, GP7/5, GP8) | SUM(GP1:3,GP7/4) | SUM(1:3,GP8,GP7/4,GP12) | SUM(GP1:3,GP7/4,GP8) |
| | Higher order oligo-mannose | The percentage of oligomannose structures from M6-M9 in total glycans | SUM(GP11, GP14/2, GP16:17, GP23:24, GP29, GP38,GP42) | SUM(GP11,GP17,GP21,GP23:24,GP29,GP42) | SUM(GP11,GP14,GP17,GP23:24,GP29) | SUM(GP11,GP16/2,GP17:18,GP23,GP25,GP29,GP42) |
| **Hybrid** | Total hybrid glycans | The percentage of hybrid structures in total glycans | SUM(GP9, GP12) | SUM(GP5,GP8,GP12, GP15) | SUM(GP8,GP12) | SUM(GP5) |
| **Core fucosy-lation** | Core $F^{Total}$ | The percentage of core fucosylated glycans in total glycans | SUM(GP2,GP4,GP7/2,GP8, GP10,GP15,GP18,GP21,GP25:26,GP31:36,GP39:41,GP44:46,GP48,GP51,GP52, GP55:56,GP58:59) | SUM(GP2,GP4,GP6,GP7/2,GP8,GP10,GP13,GP15,GP18:19,GP21,GP25/2,GP26:28,GP32,GP34,GP36,GP38,GP40:41,GP45:48,GP52,GP55:56,GP58) | SUM(GP2,GP4, GP7/2,GP8,GP10,GP13,GP15:16,GP18,GP21,GP25:28,GP34:36,GP38:41,GP43:45,GP48:49,GP52,GP55:56,GP58) | SUM(GP2,GP4,GP6/2,GP7/2,GP8,GP10,GP13:15,GP16/2,GP19,GP22,GP24,GP26,GP30,GP32,GP34:36,GP38,GP40:42,GP45:48,GP50/2,GP55:56,GP58) |
| **Outer arm fucosy-lation** | Outer arm $F^{Total}$ | The percentage of outer arm fucosylated glycans in total glycans | SUM(GP14/2,GP15, GP18:19,GP26:27,GP33:34,GP36,GP40, GP44,GP46:49,GP55,GP58:59) | SUM(GP19,GP25/2,GP26:27,GP32:34,GP36,GP39:40,GP43:44,GP46:47,GP49:51,GP54, GP56,GP58) | SUM(GP13,GP15:16,GP19,GP21,GP25:28,GP31,GP36,GP39:40,GP42:45,GP48,GP51,GP54:55,GP58:59) | SUM(GP12,GP15,GP16/2, GP19,GP21,GP24,GP26,GP27,GP34,GP36,GP38:41, GP43,GP44,GP46,GP47,GP48/2,GP49,GP50/2,GP51,GP54,GP58) |
| **N-acetyl-galacto-samine** | GalNAc | The percentage of N-acetylgalactosimined structures in total glycans | ND | SUM(GP19,GP32,GP59) | SUM(GP18,GP28,GP31,GP39,GP51,GP58) | SUM(GP9,GP12:13,GP16/2,GP19,GP41,GP52,GP59) |
| **Lac** | Total Lac structures | The percentage of structures with Lac in total glycans | SUM(GP21,GP28,GP31,GP35,GP45,GP51,GP56) | SUM(GP45,GP55) | SUM(GP34,GP47,GP56) | SUM(GP28,GP45,GP55:56) |

Table S2. Human brain tissue of Parkinson's disease (PD), incidental Lewy Bodies disease (ILBD) and control cases used in this study. Cntrl: control; PMD: post-mortem delay; N/A: not applicable

| Case number | Gender | PMD (hours) | Age at death (years) | Duration of the disease (years) | Braak stage | Brain Bank identifier number |
|---|---|---|---|---|---|---|
| **DISEASE** | | | | | | |
| PD1 | M | 21 | 79 | 12 | 3 | PD014 |
| PD2 | F | 14 | 76 | 10 | 4 | PD022 |
| PD3 | F | 10 | 80 | 13 | 4 | PD063 |
| PD4 | F | 22 | 87 | 9 | 4 | PD086 |
| PD5 | M | 9 | 72 | 6 | 4 | PD109 |
| PD6 | F | 5 | 86 | 18 | 3 | PD204 |
| PD7 | M | 11 | 72 | 2 | 3 | PD572 |
| PD8 | F | 19 | 80 | 13 | 3 | PD576 |
| PD9 | M | 9 | 76 | 21 | 4 | PD579 |
| PD10 | F | 16 | 82 | 11 | 3 | PD590 |
| PD11 | M | 17 | 77 | 9 | 4 | PD591 |
| PD12 | M | 16 | 85 | 15 | 4 | PD596 |
| PD13 | M | 11 | 78 | 16 | 4 | PD612 |
| PD14 | F | 24 | 83 | 12 | 4 | PD683 |
| PD15 | M | 16 | 86 | 19 | 3 | PD687 |
| PD16 | F | 5 | 88 | 5 | 3 | PD709 |
| PD17 | F | 17 | 87 | 13 | 4 | PD712 |
| ILBD1 | F | 10 | 104 | N/A | ILBD | PDC007 |
| ILBD2 | F | 28 | 77 | N/A | ILBD | PDC082 |
| ILBD3 | F | 39 | 82 | N/A | ILBD | PDC108 |
| **CONTROLS** | | | | | | |
| Ctrl1 | F | 17 | 71 | N/A | N/A | PDC008 |
| Ctrl2 | F | 15 | 81 | N/A | N/A | PDC015 |

| Case number | Gender | PMD (hours) | Age at death (years) | Duration of the disease (years) | Braak stage | Brain Bank identifier number |
|---|---|---|---|---|---|---|
| Ctrl3 | M | 12 | 65 | N/A | N/A | PDC022 |
| Ctrl4 | F | 23 | 80 | N/A | N/A | PDC026 |
| Ctrl5 | M | 12 | 90 | N/A | N/A | PDC034 |
| Ctrl6 | F | 15 | 61 | N/A | N/A | PDC040 |
| Ctrl7 | F | 22 | 89 | N/A | N/A | PDC053 |
| Ctrl8 | F | 20 | 82 | N/A | N/A | PDC085 |
| Ctrl9 | F | 24 | 96 | N/A | N/A | PDC088 |
| Ctrl10 | M | 11 | 95 | N/A | N/A | PDC105 |
| Ctrl11 | F | 15 | 87 | N/A | N/A | PDC107 |
| Ctrl12 | F | 23 | 88 | N/A | N/A | PDC111 |
| Ctrl13 | M | 22 | 77 | N/A | N/A | C045 |
| Ctrl14 | M | 10 | 68 | N/A | N/A | C048 |
| Ctrl15 | M | 16 | 66 | N/A | N/A | C054 |
| Ctrl16 | F | 21 | 63 | N/A | N/A | C064 |
| Ctrl17 | F | 22 | 84 | N/A | N/A | C074 |
| Ctrl18 | M | 8 | 88 | N/A | N/A | C075 |
| Ctrl19 | F | 24 | 87 | N/A | N/A | C083 |
| Ctrl20 | F | 23 | 84 | N/A | N/A | C084 |
| Ctrl21 | F | 22 | 81 | N/A | N/A | C085 |
| Ctrl22 | F | 18 | 70 | 11 | N/A | PD529 |

**Table S3. Lectins used for the microarray, binding specificity and inhibitory carbohydrate**

| Lectin | Abbreviation | Specificity | Inhibitory carbohydrate |
|---|---|---|---|
| *Concanavalin A* | ConA | α-Man, α-Glc | Mannose |
| *Narcissus Pseudonarcissus* | NPL | Terminal and internal Man | Mannose |
| *Vicia faba lectin* | VFA | α-Man, Glc, GlcNAc | Mannose |
| *Allium sativum lectin* | ASA | α-1,3Man | Mannose |
| *Galanthus nivalin agglutinin* | GNA | α-D-Man, terminal α-1,3Man | Mannose |
| *Triticum Vulgaris* | WGA | (GlcNAc)n, Sialic acid | GlcNAc |
| *Phytolacca americana* | PWA | (GlcNAc)3 | GlcNAc |
| *Griffonia simplicifolia II* | BS-II | Terminal GlcNAc | GlcNAc |
| *Lycopersicon esculentum* | LEL | (GlcNAc)3 | GlcNAc |
| *Solanum tuberosum* | STL | (GlcNAc)3, LacNAc | GlcNAc |
| *Pseudomonas aeruginosa PA-I* | PAL | Galactose | Galactose |
| *Datura stramonium* | DSL | (β-1,4) linked *N*-Acetylglucosamine oligomers, (GlcNAc)2-3, LacNAc | GlcNAc |
| *Ricinum communis agglutinin* | $RCA_{120}$ | β -Gal, Lac, LacNAc | Galactose |
| *Erythrina cristagalli A* | ECA | Gal, GalNAc | GalNAc |
| *Phaseolus vulgaris agglutinin* | PHA E+L | Oligosaccharides | Lactose |
| *Griffonia simplicifolia I* | BS-I-B4 | α-Gal | Galactose |
| *Cicer arietinum* | CAL | Fetuin, Lac, IgM | Lactose |
| *Maackia amurensis Lectin II* | MAL-II | LacNAc | Galactose |
| *Sambucus nigra* | SNA | Sialic acid-α-2,6–GalNAc | Lactose, Sialic acid |
| *Maackia amurensis Lectin I* | MAL-I | Sialic acid-α-2,3–GalNAc | Lactose, Sialic acid |
| *Homarus americanus* | HMA | LAg1 NeuNAc. LAg2: GalNAc | GalNAc |
| *Ulex Europaea Agglutinin* | UEA-I | α-1,2-fucose | Fucose |
| *Pisum sativum* | PSA | Fucose -α-1,6-GlcNAc, α-Man | Fucose |
| *Lotus tetragonolobus* | LTL | Terminal α-Fucose, $Le^x$ | Fucose |
| *Aspergillus orzyae* | AOL | Fucose | Fucose |
| *Aleuria aurantia lectin* | AAL | α –1,6-Fuc to *N*-Acetylglucosamine, | Fucose |

| | | | |
|---|---|---|---|
| | | α-1,3-Fuc to *N*-Acetyllactosamine | |
| *Anguilla anguilla agglutinin* | AAA | D-Fucose, α-1,2-Fucose, α-1,4-Fucose | Fucose |
| *Vicia villosa B4* | VVL | GalNAc | GalNAc |
| *Jacalin* | JAC | T antigen | GalNAc |
| *Agaricus bisporus* | ABL | Gal- β 1,3-GalNAc, T antigen | Galactose |
| *Amarantus Caudatus* | ACA | T antigen, (Gal-β 1,3-GalNAc α -Thr/Ser) | Galactose |
| *Maclura Pomifera* | MPA | T antigen, αGalNAc | Galactose |
| *Psophocarpus tetragonolobus II* | PT-II | α-1,2-fucosylated LacNAc | Galactose |
| *Griffonia simplicifolia I* | BS-I | α-Gal | Galactose |
| *Euonymus Europaeus* | EEA | Lac, blood groups B and H | Galactose |
| *Marasmium oreades agglutinin* | MOA | Gal-α1,3Gal and Gal-α1,3Gal-β1,4GlcNAc | Galactose |
| *Helix pomatia* | HPL | α-GalNAc terminal | GalNAc |
| *Sophora japonica* | SJA | GalNAc | GalNAc |
| *Helix aspersa lectin* | HAL | α-GalNAc terminal | GalNAc |
| *Soybean agglutinin* | SBA | α-Gal-GalNAc | GalNAc |
| *Peanut agglutinin* | PNA | T antigen, Gal β-1,3-GalNAc | Galactose |
| *Dolichos biflorus agglutinin* | DBA | Terminal GalNAc | GalNAc |
| *Wisteria floribunda* | WFL | GalNAc | GalNAc |
| *Salvia sclarea lectin* | SSA | Terminal GalNAc linked to serine | GalNAc |
| *Psophocarpus tetragonolobus I* | PT-I | α-GalNAc | GalNAc |
| *Bauhinia Purpurea lectin* | BPL | GalNAc | GalNAc |

**Table S4. Lectins used for cytochemistry, their binding specificity, concentration used and inhibitory carbohydrate (haptenic sugar, 150 mM).**

| Lectin | Abbreviation | Binding specificity | Concentration | Inhibitory carbohydrate |
|---|---|---|---|---|
| *Sambucus nigra agglutinin isolectin*-I | SNA-I | Neu5Ac/Gc-α(2,6)-Gal/GalNAc-R | 20 µg/mL | Lactose |
| *Anguilla anguilla agglutinin* | AAA | D-Fucose, α-1,2-Fucose, α-1,4-Fucose | 10 µg/mL | Fucose |
| *Datura stramonium* | DSL | (β-1,4) linked N-acetylglucosamine oligomers, (GlcNAc)2-3, LacNAc | 15 µg/mL | GlcNAc, GalNAc |
| *Ricinus communis* Agglutinin | RCA | β -Gal, Lac, LacNAc | 5 µg/mL | Gal or lactose |
| *Galanthus nivalin agglutinin* | GNA | α-D-Man, terminal α-1,3Man | 10 µg/mL | Mannose |

**Table S5. Genes present in the PCR RT2- Profiler Human Glycosylation Array (Qiagen, UK)**

| RefSeq Number | Description | Symbol |
|---|---|---|
| NM_016161 | Alpha-1,4-N-acetylglucosaminyltransferase | A4GNT |
| NM_000027 | Aspartylglucosaminidase | AGA |
| NM_194318 | Beta 1,3-galactosyltransferase-like | B3GLCT |
| NM_006577 | UDP-GlcNAc:betaGal beta-1,3-N-acetylglucosaminyltransferase 2 | B3GNT2 |
| NM_014256 | UDP-GlcNAc:betaGal beta-1,3-N-acetylglucosaminyltransferase 3 | B3GNT3 |
| NM_030765 | UDP-GlcNAc:betaGal beta-1,3-N-acetylglucosaminyltransferase 4 | B3GNT4 |
| NM_198540 | UDP-GlcNAc:betaGal beta-1,3-N-acetylglucosaminyltransferase 8 | B3GNT8 |
| NM_001497 | UDP-Gal:betaGlcNAc beta 1,4- galactosyltransferase, polypeptide 1 | B4GALT1 |
| NM_003780 | UDP-Gal:betaGlcNAc beta 1,4- galactosyltransferase, polypeptide 2 | B4GALT2 |
| NM_003779 | UDP-Gal:betaGlcNAc beta 1,4- galactosyltransferase, polypeptide 3 | B4GALT3 |
| NM_004776 | UDP-Gal:betaGlcNAc beta 1,4- galactosyltransferase, polypeptide 5 | B4GALT5 |
| NM_020156 | Core 1 synthase, glycoprotein-N-acetylgalactosamine 3-beta-galactosyltransferase, 1 | C1GALT1 |
| NM_152692 | C1GALT1-specific chaperone 1 | C1GALT1C1 |
| NM_014674 | ER degradation enhancer, mannosidase alpha-like 1 | EDEM1 |
| NM_018217 | ER degradation enhancer, mannosidase alpha-like 2 | EDEM2 |
| NM_025191 | ER degradation enhancer, mannosidase alpha-like 3 | EDEM3 |
| NM_000147 | Fucosidase, alpha-L- 1, tissue | FUCA1 |
| NM_032020 | Fucosidase, alpha-L- 2, plasma | FUCA2 |
| NM_173540 | Fucosyltransferase 11 (alpha (1,3) fucosyltransferase) | FUT11 |
| NM_178157 | Fucosyltransferase 8 (alpha (1,6) fucosyltransferase) | FUT8 |
| NM_020474 | UDP-N-acetyl-alpha-D-galactosamine:polypeptide N-acetylgalactosaminyltransferase 1 (GalNAc-T1) | GALNT1 |
| NM_198321 | UDP-N-acetyl-alpha-D-galactosamine:polypeptide N-acetylgalactosaminyltransferase 10 (GalNAc-T10) | GALNT10 |
| NM_022087 | UDP-N-acetyl-alpha-D-galactosamine:polypeptide N-acetylgalactosaminyltransferase 11 (GalNAc-T11) | GALNT11 |
| NM_024642 | UDP-N-acetyl-alpha-D-galactosamine:polypeptide N-acetylgalactosaminyltransferase 12 (GalNAc-T12) | GALNT12 |
| NM_052917 | UDP-N-acetyl-alpha-D-galactosamine:polypeptide N-acetylgalactosaminyltransferase 13 (GalNAc-T13) | GALNT13 |

| RefSeq Number | Description | Symbol |
|---|---|---|
| NM_024572 | UDP-N-acetyl-alpha-D-galactosamine:polypeptide N-acetylgalactosaminyltransferase 14 (GalNAc-T14) | GALNT14 |
| NM_004481 | UDP-N-acetyl-alpha-D-galactosamine:polypeptide N-acetylgalactosaminyltransferase 2 (GalNAc-T2) | GALNT2 |
| NM_004482 | UDP-N-acetyl-alpha-D-galactosamine:polypeptide N-acetylgalactosaminyltransferase 3 (GalNAc-T3) | GALNT3 |
| NM_003774 | UDP-N-acetyl-alpha-D-galactosamine:polypeptide N-acetylgalactosaminyltransferase 4 (GalNAc-T4) | GALNT4 |
| NM_007210 | UDP-N-acetyl-alpha-D-galactosamine:polypeptide N-acetylgalactosaminyltransferase 6 (GalNAc-T6) | GALNT6 |
| NM_017423 | UDP-N-acetyl-alpha-D-galactosamine:polypeptide N-acetylgalactosaminyltransferase 7 (GalNAc-T7) | GALNT7 |
| NM_017417 | UDP-N-acetyl-alpha-D-galactosamine:polypeptide N-acetylgalactosaminyltransferase 8 (GalNAc-T8) | GALNT8 |
| NM_021808 | UDP-N-acetyl-alpha-D-galactosamine:polypeptide N-acetylgalactosaminyltransferase 9 (GalNAc-T9) | GALNT9 |
| NM_020692 | UDP-N-acetyl-alpha-D-galactosamine:polypeptide N-acetylgalactosaminyltransferase-like 1 | GALNT16 |
| NM_145292 | UDP-N-acetyl-alpha-D-galactosamine:polypeptide N-acetylgalactosaminyltransferase-like 5 | GALNTL5 |
| NM_001034845 | UDP-N-acetyl-alpha-D-galactosamine:polypeptide N-acetylgalactosaminyltransferase-like 6 | GALNTL6 |
| NM_198334 | Glucosidase, alpha; neutral AB | GANAB |
| NM_001490 | Glucosaminyl (N-acetyl) transferase 1, core 2 | GCNT1 |
| NM_004751 | Glucosaminyl (N-acetyl) transferase 3, mucin type | GCNT3 |
| NM_016591 | Glucosaminyl (N-acetyl) transferase 4, core 2 | GCNT4 |
| NM_000404 | Galactosidase, beta 1 | GLB1 |
| NM_024312 | N-acetylglucosamine-1-phosphate transferase, alpha and beta subunits | GNPTAB |
| NM_032520 | N-acetylglucosamine-1-phosphate transferase, gamma subunit | GNPTG |
| NM_000520 | Hexosaminidase A (alpha polypeptide) | HEXA |
| NM_000521 | Hexosaminidase B (beta polypeptide) | HEXB |
| NM_005907 | Mannosidase, alpha, class 1A, member 1 | MAN1A1 |
| NM_006699 | Mannosidase, alpha, class 1A, member 2 | MAN1A2 |
| NM_016219 | Mannosidase, alpha, class 1B, member 1 | MAN1B1 |
| NM_020379 | Mannosidase, alpha, class 1C, member 1 | MAN1C1 |

| RefSeq Number | Description | Symbol |
|---|---|---|
| NM_002372 | Mannosidase, alpha, class 2A, member 1 | MAN2A1 |
| NM_006122 | Mannosidase, alpha, class 2A, member 2 | MAN2A2 |
| NM_000528 | Mannosidase, alpha, class 2B, member 1 | MAN2B1 |
| NM_005908 | Mannosidase, beta A, lysosomal | MANBA |
| NM_002406 | Mannosyl (alpha-1,3-)-glycoprotein beta-1,2-N-acetylglucosaminyltransferase | MGAT1 |
| NM_002408 | Mannosyl (alpha-1,6-)-glycoprotein beta-1,2-N-acetylglucosaminyltransferase | MGAT2 |
| NM_002409 | Mannosyl (beta-1,4-)-glycoprotein beta-1,4-N-acetylglucosaminyltransferase | MGAT3 |
| NM_012214 | Mannosyl (alpha-1,3-)-glycoprotein beta-1,4-N-acetylglucosaminyltransferase, isozyme A | MGAT4A |
| NM_014275 | Mannosyl (alpha-1,3-)-glycoprotein beta-1,4-N-acetylglucosaminyltransferase, isozyme B | MGAT4B |
| NM_013244 | Mannosyl (alpha-1,3-)-glycoprotein beta-1,4-N-acetylglucosaminyltransferase, isozyme C (putative) | MGAT4C |
| NM_002410 | Mannosyl (alpha-1,6-)-glycoprotein beta-1,6-N-acetyl-glucosaminyltransferase | MGAT5 |
| NM_144677 | Mannosyl (alpha-1,6-)-glycoprotein beta-1,6-N-acetyl-glucosaminyltransferase, isozyme B | MGAT5B |
| NM_006302 | Mannosyl-oligosaccharide glucosidase | MOGS |
| NM_016256 | N-acetylglucosamine-1-phosphodiester alpha-N-acetylglucosaminidase | NAGPA |
| NM_000434 | Sialidase 1 (lysosomal sialidase) | NEU1 |
| NM_005383 | Sialidase 2 (cytosolic sialidase) | NEU2 |
| NM_006656 | Sialidase 3 (membrane sialidase) | NEU3 |
| NM_080741 | Sialidase 4 | NEU4 |
| NM_181673 | O-linked N-acetylglucosamine (GlcNAc) transferase (UDP-N-acetylglucosamine:polypeptide-N-acetylglucosaminyl transferase) | OGT |
| NM_172236 | Protein O-fucosyltransferase 1 | POFUT1 |
| NM_133635 | Protein O-fucosyltransferase 2 | POFUT2 |
| NM_017739 | Protein O-linked mannose beta1,2-N-acetylglucosaminyltransferase | POMGNT1 |
| NM_007171 | Protein-O-mannosyltransferase 1 | POMT1 |
| NM_013382 | Protein-O-mannosyltransferase 2 | POMT2 |
| NM_002743 | Protein kinase C substrate 80K-H | PRKCSH |

| RefSeq Number | Description | Symbol |
|---|---|---|
| NM_173344 | ST3 beta-galactoside alpha-2,3-sialyltransferase 1 | ST3GAL1 |
| NM_006927 | ST3 beta-galactoside alpha-2,3-sialyltransferase 2 | ST3GAL2 |
| NM_003032 | ST6 beta-galactosamide alpha-2,6-sialyltranferase 1 | ST6GAL1 |
| NM_018414 | ST6 (alpha-N-acetyl-neuraminyl-2,3-beta-galactosyl-1,3)-N-acetylgalactosaminide alpha-2,6-sialyltransferase 1 | ST6GALNAC1 |
| NM_006011 | ST8 alpha-N-acetyl-neuraminide alpha-2,8-sialyltransferase 2 | ST8SIA2 |
| NM_015879 | ST8 alpha-N-acetyl-neuraminide alpha-2,8-sialyltransferase 3 | ST8SIA3 |
| NM_175052 | ST8 alpha-N-acetyl-neuraminide alpha-2,8-sialyltransferase 4 | ST8SIA4 |
| NM_001004470 | ST8 alpha-N-acetyl-neuraminide alpha-2,8-sialyltransferase 6 | ST8SIA6 |
| NM_020120 | UDP-glucose glycoprotein glucosyltransferase 1 | UGGT1 |
| NM_020121 | UDP-glucose glycoprotein glucosyltransferase 2 | UGGT2 |
| NM_001101 | Actin, beta | ACTB |
| NM_004048 | Beta-2-microglobulin | B2M |
| NM_002046 | Glyceraldehyde-3-phosphate dehydrogenase | GAPDH |
| NM_000194 | Hypoxanthine phosphoribosyltransferase 1 | HPRT1 |
| NM_001002 | Ribosomal protein, large, P0 | RPLP0 |

**Table S6. List of primers used for RT-PCR.**

| Gene/Primer | Forward sequence | Reverse sequence |
|---|---|---|
| 18sc (housekeeping) | AATCAGTTATGGTTCCTTTGTCG | GCTCTAGAATTACCACAGTTATCCAA |
| ATF6 | GGACCAGGTGGTGTCAGAG | GACAGCTCTGCGCTTTGG |
| CHOP | GAAATCGAGCGCCTGACCAG | GGAGGTGATGCCAACAGTTCA |
| XBP1 unspliced | CAGACTACGTGCGCCTCTG | CTTCTGGGTAGACCTCTGGG |
| XBP1 spliced | GAGTCCGCAGCAGGTGC | GGTCCAACTTGTCCAGAATGC |

**Table S7 (Datasheet). Exoglycosidase digestion panels for healthy striatum pool (Tables S7.1/S7.2), PD striatum pool (Tables S7.3/S7.4), healthy substantia nigra pool (Tables S7.5/S7.6), PD substantia nigra pool (Tables S7.7/S7.8). –** REFER TO SUPPLEMENTARY SPREADSHEET FILE.

**Table S8 (Datasheet). HILIC-UPLC-MS data from healthy striatum pool (Table S8.1), PD striatum pool (Table S8.2), healthy substantia nigra pool (Table S8.3), PD substantia nigra pool (Table S8.4), outlining m/z values as well as elution time (min), relative charge state observed, error (ppm) and monosaccharide composition. -** REFER TO SUPPLEMENTARY SPREADSHEET FILE.
